# A synthetic small molecule stalls pre-mRNA splicing by promoting an early-stage U2AF2–RNA complex

**DOI:** 10.1101/2020.09.28.317727

**Authors:** Rakesh Chatrikhi, Callen F. Feeney, Mary J. Pulvino, Georgios Alachouzos, Andrew J. MacRae, Zackary Falls, Sumit Rai, William W. Brennessel, Jermaine L. Jenkins, Matthew J. Walter, Timothy A. Graubert, Ram Samudrala, Melissa S. Jurica, Alison J. Frontier, Clara L. Kielkopf

## Abstract

Dysregulated pre-mRNA splicing is an emerging Achilles heel of cancers and myelodysplasias. To expand the currently limited portfolio of small molecule drug leads, we screened for chemical modulators of the U2AF complex, which nucleates spliceosome assembly and is mutated in myelodysplasias. A hit compound specifically enhances RNA binding by a U2AF2 subunit. Remarkably, the compound inhibits splicing of representative substrates in cells and stalls spliceosome assembly at the stage of U2AF function. Computational docking, together with structure-guided mutagenesis, indicates that the compound bridges an active conformation of the U2AF2 RNA recognition motifs *via* hydrophobic and electrostatic moieties. Altogether, our results highlight the potential of trapping early spliceosome assembly as an effective pharmacological means to manipulate pre-mRNA splicing. By extension, we suggest that stabilizing inactive checkpoints may offer a breakthrough approach for small molecule inhibition of multi-stage macromolecular assemblies.

## Introduction

Alternative splicing of human pre-mRNAs is an essential step of gene expression and generates a multitude of tissue-specific isoforms from a much smaller number of genes^1,2^. Genomes of hematologic malignances acquire recurrent mutations among a subset of pre-mRNA splicing factors (U2AF1, SF3B1, SRSF2, ZRSR2) (reviewed in^3^). The mutational hotspots typically cluster at key interfaces involved in the early stages of spliceosome assembly^4^. Dysregulated pre-mRNA splicing resulting from these mutations combined with other transcriptional abnormalities renders hematologic malignancies and cancers more sensitive to spliceosome inhibitors relative to their “normal” counterparts^5–12^. As such, defective pre-mRNA splicing offers a promising target for potential anti-cancer therapies (reviewed in^13^). The most clinically-advanced inhibitors of pre-mRNA splicing include small molecules targeting a druggable site of SF3B1 adjacent a conserved pre-mRNA adenosine (branchpoint)^12,14,15^. These SF3B1-targeted family of inhibitors selectively arrest growth of cancer cells in a variety of models and patient samples^5–12,16^.

In view of the large uncharted pharmacological space created by the complexity of the spliceosome, there is rich promise for compounds that target any of the more than a hundred splicing factors beyond SF3B1. One promising splicing factor, for which the potential as a therapeutic target has yet to be explored, is the U2 small nuclear ribonucleoprotein auxiliary factor (U2AF). A heterodimer of U2AF2 and U2AF1 subunits recognizes and nucleates spliceosome assembly at the polypyrimidine (Py) tract^17^ and AG-dinucleotide^18,19^ consensus signals of the 3’ splice site. U2AF initially binds as a ternary complex with a third splicing factor, SF1, which subsequently is displaced by the SF3B1 subunit of the U2 particle. U2AF2 supports splicing at most sites, but depends on assistance from the U2AF1 subunit when the Py tract signal is degenerate^18,19^. Notably, the U2AF1 subunit frequently carries an S34F/Y mutation, or less frequently Q157R/P, in approximately 12% of patients with myelodysplastic syndromes (MDS)^20,21^ or 3% with lung adenocarcinomas^22^. Interestingly, U2AF1 is a paralogue of ZRSR2, another recurrently mutated splicing factor in MDS and chronic myelomonocytic leukemia, which functions as part of the minor U12 rather than U2-type spliceosome machinery^23,24^. Albeit at lower frequencies, the U2AF2 subunit also acquires mutations in cancers and leukemias^25–27^. Since U2AF2 recruits the SF3B1 subunit, the pre-clinical success of SF3B1 inhibitors suggests that targeting U2AF2 likewise could produce small molecule tools for investigating spliceosome assembly in normal cells and hematologic malignancies.

Here, we identify and characterize a small molecule modulator (NSC 194308) that increased, rather than reduced, association of the U2AF1–U2AF2–SF1–splice site RNA complex by binding a site between the U2AF2 RNA recognition motifs (RRM1 and RRM2). NSC 194308 inhibited pre-mRNA splicing by stalling spliceosome assembly at the point where U2AF helps recruit U2 snRNP to the branchpoint. These results set the stage for future optimization of a promising hit inhibitor of pre-mRNA splicing, and highlight a general strategy for inhibiting multi-stage processes by stabilizing early checkpoints.

## Results

### Establishing a Screen for U2AF–RNA Modulators

To screen for small molecule modulators of the U2AF-containing spliceosome, we focused on a complex among the U2AF1, U2AF2, and SF1 splicing factors (“U2AF”-complex) that recognizes the 3’ splice site in the initial stage of spliceosome assembly. Our U2AF protein complex included the relevant RNA binding domains (**Fig. 1a**) and was expressed and purified as described^28,29^. Fluorescence polarization (FP) of a 5’-fluorescein-labeled RNA oligonucleotide (^Fl^RNA) was the principle readout of the screen (**Fig. 1b**). Association of the ribonucleoprotein slows rotational diffusion of the ^Fl^RNA and hence retained more of the anisotropy of fluorophore emission following excitation with polarized light. In addition to comprising the relevant complex for the early stage of 3’ splice site recognition, the inclusion of all three splicing factor subunits is expected to improve the signal of the screen by increasing the change in FP following ^Fl^RNA binding to a greater extent than a single subunit alone. We included the MDS-associated S34F mutation of the U2AF1 subunit as a potential disease-relevant feature for recognition by small molecules. Likewise, we chose variants of an S34F-affected splice site of the *DEK* oncogene^28^ as the RNA binding site of the targeted U2AF complex in the screen, specifically a *DEK(-3U)* ^Fl^RNA site for the U2AF – RNA enhancer screen. We screened the 1593-compound Diversity Set V of the National Cancer Institute (NCI) Developmental Therapeutics Program, for which the modest size was suitable considering the yields of our protein preparation and provided a strong starting point of diverse, pharmacologically-desirable features for subsequent optimization of initial hits. Our initial search for inhibitors of binding resulted in three aromatic compounds that disrupted U2AF – RNA complexes (**Supplementary Results, Supplementary Table 1, Supplementary Fig. 1a-e**). Rather than the U2AF – RNA complex, these candidate inhibitors appeared to target the RNA by nonspecific intercalation (**Supplementary Results, Supplementary Fig. 1f**). Instead, we prioritized the provocative results of a screen for small molecule enhancers of U2AF–RNA association, as described below.

**Fig. 1.**
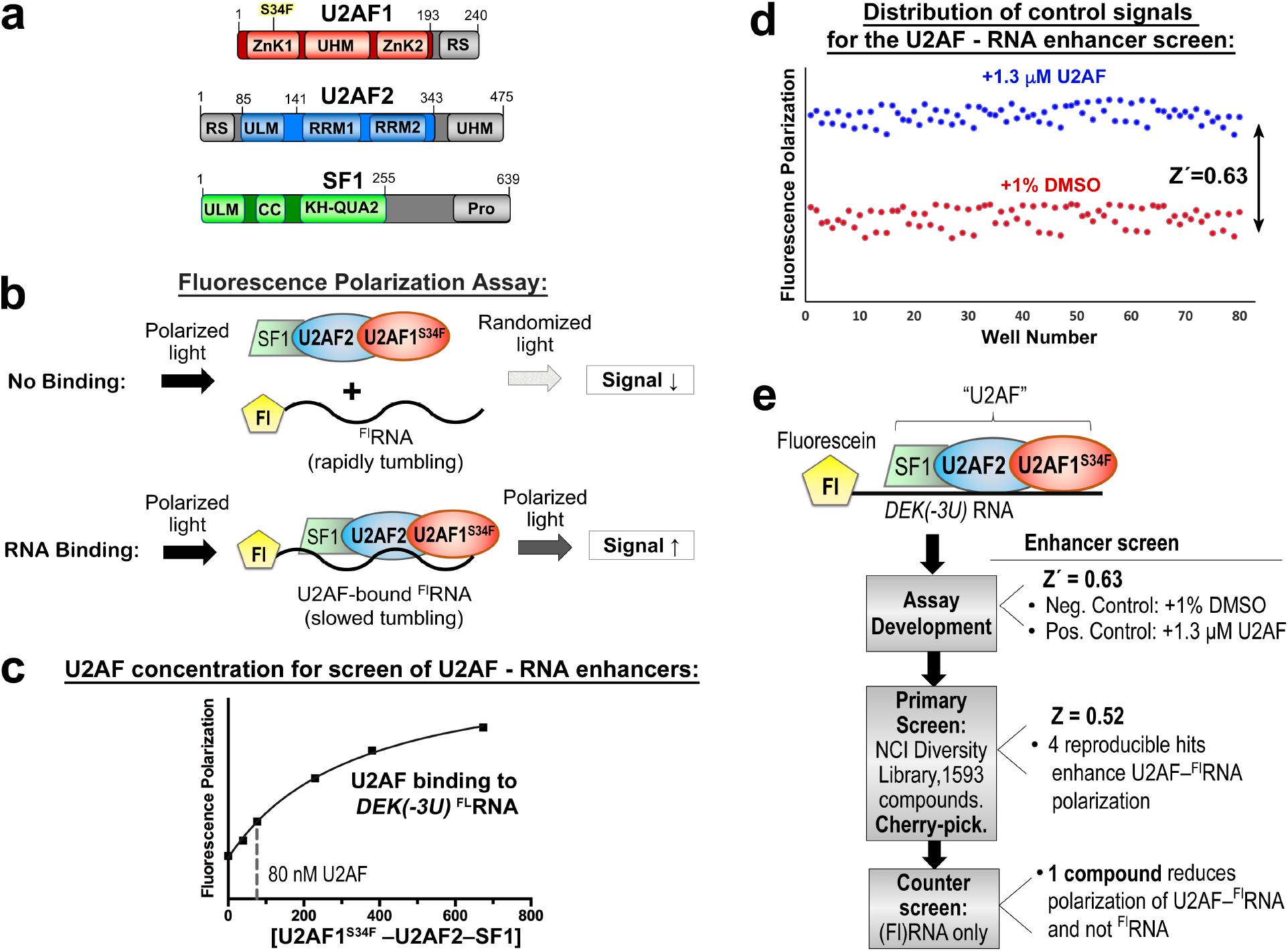
Implementation of high-throughput screen for modulators of U2AF–RNA complexes. (**a**) Domains of U2AF1, U2AF2, and SF1 splicing factors. Gray regions were excluded from the expression constructs used for the screen. (**b**) Schematic diagram of the fluorescence polarization RNA binding assay. Fl, 5' fluorescein. (**c**) Fluorescence polarization binding curve for U2AF1^S34F^–U2AF2–SF1 titrated into fluorescein *DEK(-3U)* ^FL^RNA. The nonlinear fits are overlaid on the average data points of three replicates as a solid curve. A dashed line marks the protein concentration used in each screen. (**d**) Distribution of positive (blue) and negative (red) control signals used for the calculation of the Z’-factors of the assay. (**e**) Overview of the screens for enhancers (results to right).

### Results of a High-Throughput Screen for U2AF–RNA Enhancers

Before screening chemical libraries, we first assessed the quality of the assay by measuring the FP of a plate comprising alternating rows of positive and negative control samples. We chose a baseline concentration of U2AF1^S34F^–U2AF2–SF1 proteins (80 nM) within the initial phase of the *DEK(-3U)* ^Fl^RNA binding curve (**Fig. 1c**). The effective solvent of the compounds (1% v/v DMSO) served as a negative control. As a positive control for enhancers of protein–RNA binding, we added excess U2AF1^S34F^–U2AF2–SF1 protein in a buffer including DMSO (1% v/v) to match the compound-containing samples. We reproducibly found that addition of unlabeled RNA reduced the FP, excess U2AF protein increased FP, and the trivial amount of DMSO solvent had no detectable effect on the binding reaction. From these distributions, we calculated a statistical Z’-factor to evaluate the quality of the assay (**Fig. 1d**). The Z’-factor^30^ of 0.63 for the set-up of the enhancer screens convincingly showed that the FP assay could measure gain of U2AF–RNA binding and set the stage to proceed with a chemical library screen.

We next explored the possibility that small molecules could increase U2AF binding to weak splice sites. We screened the NCI Diversity Set V library for enhancers of the U2AF1^S34F^–U2AF2– SF1–*DEK(-3U)* ^Fl^RNA complex (**Fig. 1c, Supplementary Table 1**). The Z-factor of the assay was 0.52, indicating that hits could be expected under the conditions of this screen^30^. An initial pool of seven compounds (0.4% of the starting library) increased FP by at least 75% compared to a positive control signal (excess U2AF1^S34F^–U2AF2–SF1 protein). Replicates showed that four of these compounds (0.25% of the starting library) reproducibly increased the FP of the U2AF–RNA complex by >75% compared to the positive control. However, only a single compound (NSC 194308) specifically increased the FP of the U2AF1^S34F^–U2AF2–SF1–*DEK(-3U)* ^Fl^RNA ribonucleoprotein without affecting the *DEK(-3U)* ^Fl^RNA alone. The candidate U2AF–RNA enhancer lacked the aromatic rings of the inhibitors identified in our first screen, and instead included distinct aliphatic bicarbocyclic and zwitterionic groups (**Fig. 2a,** inset). The concentration of half-maximal enhancement (EC_50_ ~100 μM) was estimated from the increased FP of the U2AF1^S34F^–U2AF2–SF1–*DEK(-3U)* ^Fl^RNA complex during titration with the hit enhancer (**Fig. 2a**). Although the potency of the compound was low in this assay with the purified protein complex, we viewed the specific, ribonucleoprotein-directed activity of NSC 194308 as a promising start point for further investigation and optimization, particularly considering that small perturbations of the essential U2AF complex could have potentially significant consequences for living cells.

**Fig. 2.**
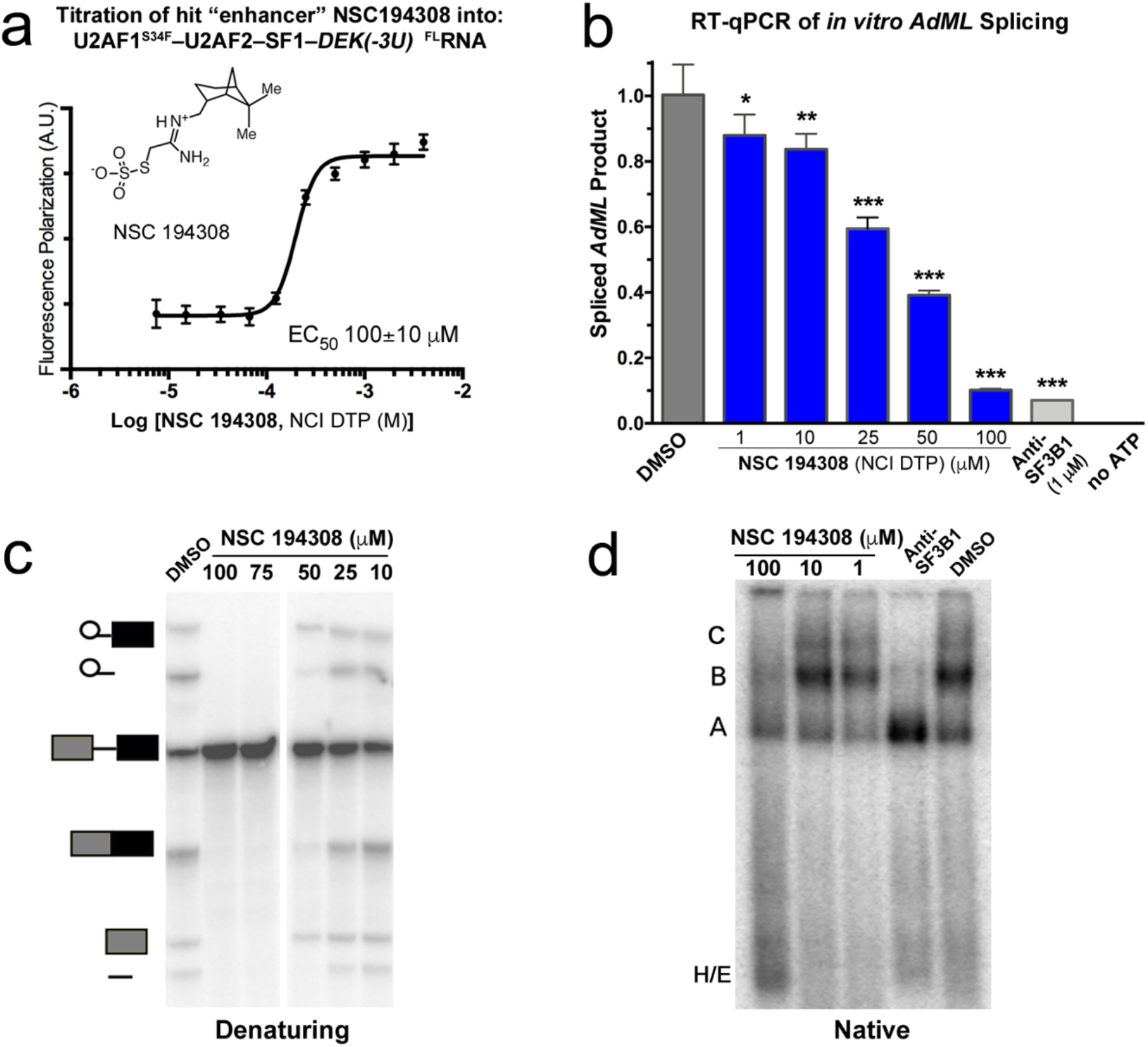
The hit enhancer of U2AF2 – RNA complexes stalls *in vitro* pre-mRNA splicing in HeLa nuclear extract at the H/E-stage. (**a**) FP dose-response of the hit enhancer (NSC194308, chemical structure inset) titrated into U2AF1^S34F^–U2AF2–SF1–*DEK(-3U)* ^FL^RNA. Data points and error bars represent the mean and standard deviations of three experimental replicates. (**b**) RT-qPCR of spliced *AdML* products. The relative amounts of spliced product were normalized to a mock-treated negative control (1% v/v DMSO, gray).Two-tailed unpaired t-tests with Welch’s correction calculated in GraphPad Prism compared the indicated NSC 194308 concentration with the mock-treated control: *, *p*<0.05; **, *p*<0.005; ***, *p*<0.0005. The positive control is an SF3B1 inhibitor (“anti-SF3B1”, 1 μM pladienolide-B). (**c**) Denaturing gel analysis of radiolabeled pre-mRNA substrate and spliced products from reactions treated with the indicated concentrations of NSC 194308 or mock-treated control. Spliced products are schematically diagrammed to the left. (**d**) Native gel analysis of spliceosome assembly at 30 minutes in the presence of the indicated concentrations of NSC 194308, positive control (“anti-SF3B1”, 1 μM spliceostatin-A), or mock-treated control. The identity of spliceosome complexes is indicated to the left with assembly occurring in the following order: H/E → A → B → C. NCI DTP, NCI Developmental Therapeutics Program.

### Chemical synthesis of NSC 194308 hit compound

To obtain sufficient, pure NSC 194308 for the investigation of its functional activities, we synthesized the compound starting from (-)-*cis*-myrtanylamine (Online Methods). Following sequential reactions with chloroacetonitrile under strongly basic conditions followed by displacement of the chloride with sodium thiosulfate (**Scheme 1**), the compound was precipitated by acidification and its identity confirmed by NMR and X-ray crystallography of the intermediate (**Compound 2** in **Scheme 1,** Online Methods) and NMR of the final product (**Compound 1** in **Scheme 1,** Online Methods) (**Supplementary Fig. 2a-b**). The product showed similar potency for inhibition of *in vitro* pre-mRNA splicing as the compound obtained from the NCI repository (**Supplementary Fig. 2c** compared to **Fig. 2b**). We also synthesized a (-)-*cis*-myrtanyl acetamidinium (MAA) variant (**Compound 3** in **Scheme 2**) to probe structure-activity relationships (described below) and confirmed its identity by NMR and X-ray crystallography (**Supplementary Fig. 3a-b**). Crystal structures of the NSC 194308 intermediate (**Supplementary Fig. 2a**) and the MAA variant (**Supplementary Fig. 3a**) established the absolute stereochemistry of the hit compound. We synthesized NSC 194308 *de novo* for use in subsequent experiments that required large amounts of material.

### Hit U2AF–RNA “Enhancer” NSC 194308 blocks the pre-mRNA splicing process

To test pre-mRNA splicing activities without potential complications from the intracellular metabolism or transport of the hit compound, we first examined the efficiencies of *in vitro* splicing reactions in HeLa nuclear extracts treated with increasing concentrations of NSC 194308 (**Fig. 2b-c**). We used quantitative real-time reverse transcription (qRT)-PCR to detect splicing of a prototypical strong splice site from the Adenovirus Major Late promoter transcript (*AdML*). We compared a known SF3B1 inhibitor (pladienolide B) alongside mock-treatment with an equivalent amount of solvent (1% v/v DMSO) as positive and negative controls. Contrary to our notion that an enhancer of U2AF–RNA interaction would stimulate pre-mRNA splicing, the NSC 194308 hit reduced splicing of the *AdML* substrate in the nuclear extract. We confirmed this result by denaturing gel electrophoresis to visualize the intermediates and products resulting from *in vitro* splicing of a radiolabeled pre-mRNA substrate (an *AdML* variant, **Fig. 2c**). The pre-mRNA substrate appeared intact at the highest concentrations of NSC 194308, which ruled out nonspecific RNA degradation due to the compound. We concluded that the apparent concentration of half-maximal inhibition of *in vitro* splicing by NSC 194308 (IC_50_ ~35 μM, **Fig. 2b-c**, **Supplementary Fig. 2c**) was similar in magnitude to its enhancement of the purified U2AF-containing ribonucleoprotein (**Fig. 2a** and **3e**).

**Fig. 3.**
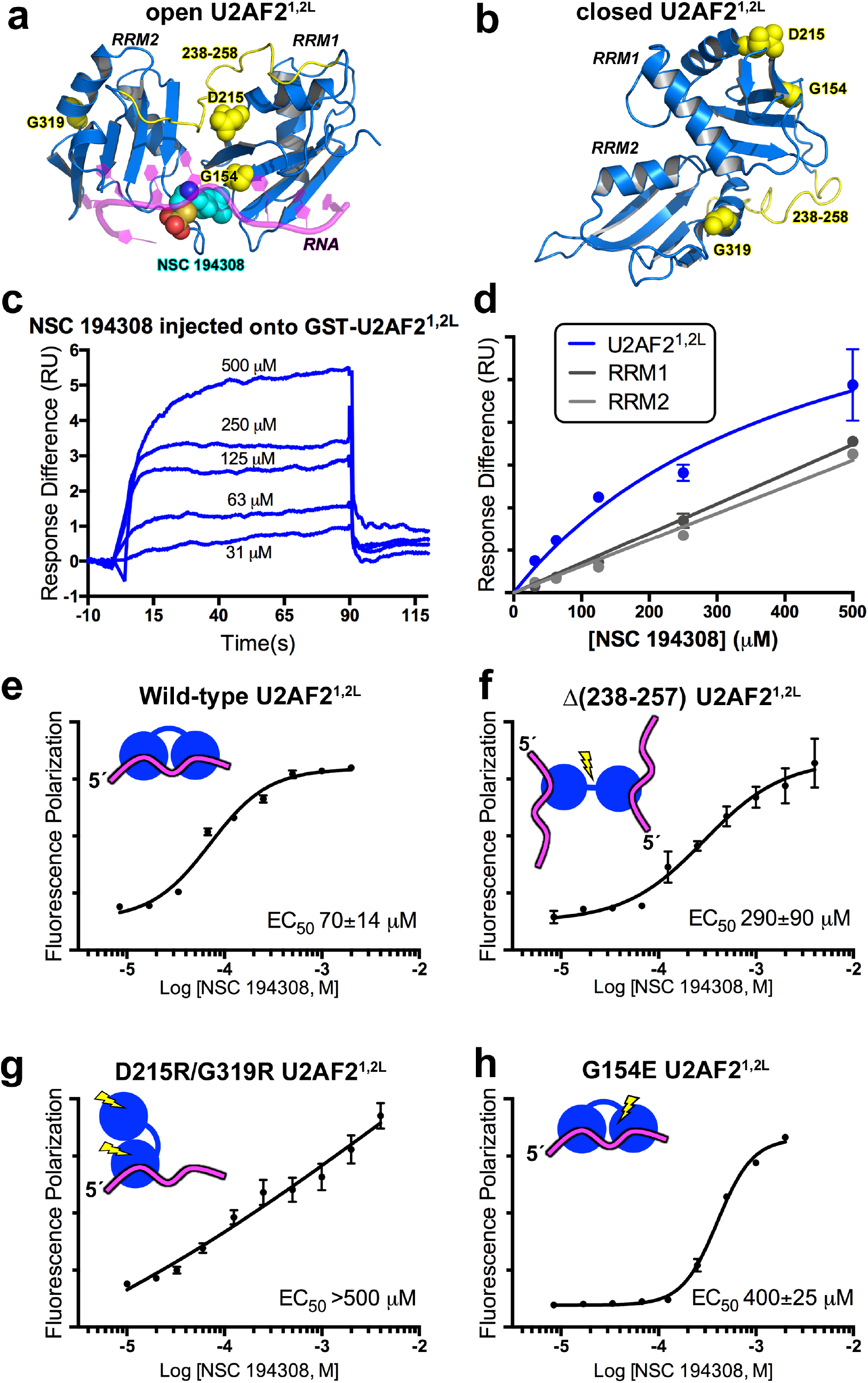
The hit enhancer NSC 194308 binds between the tandem U2AF2 RRM1/RRM2 (U2AF2^1,2L^). (**a**) The favorable predicted binding site for NSC 194308 is located between RRM1 and RRM2 (blue) of the open U2AF2^1,2L^ conformation (PDB ID 5EV4). The Py tract oligonucleotide (magenta) of the structure is overlaid as a semi-transparent cartoon for reference. Mutated regions tested in FP assays are colored yellow and labeled; site-specific mutated residues are shown as space-filling CPKs; NSC 194308 is a CPK representation colored by atom: carbon, cyan; oxygen, red; sulfur, orange; nitrogen, blue) (**b**) Locations of the mutated-regions (yellow) shown on the NMR model (blue) of closed U2AF2^1,2L^ (PDB ID 2YH0). (**c**) Representative sensorgram showing the aligned responses of a series of NSC 194308 injections at the indicated concentration onto an immobilized GST- U2AF2^1,2L^ surface. (**d**) Plot of the responses, averaged from the saturated regions of the sensorgrams in two independent replicates and fit to a nonlinear steady-state model, are shown for NSC 194308 binding to GST- U2AF2^1,2L^ (blue) or the separate, GST-RRM1 (dark gray) or GST-RRM2 (light gray). (**e-h**) FP dose-responses of the hit enhancer titrated into *DEK(-3U)* ^FL^RNA bound to U2AF2^1,2L^ variants, including (**e**) wild-type, (**f**) inter-RRM linker deletion, (**g**) D215R/G319R, or (**h**) G154E. The average and standard deviations of three experimental replicates are shown. Schematic diagrams are inset of the expected conformations of the U2AF2 mutants with blue RRMs, yellow mutations, and magenta RNA strand.

To determine the mechanism of action for NSC 194308 inhibition of splicing, we proceeded to analyze the intermediates of spliceosome assembly in the presence of increasing concentrations of the compound using native gel electrophoresis (**Fig. 2d**). The spliceosome assembles through a series of conformational and compositional changes that sequentially form the H-, E-, A-, B- and C-complexes (reviewed in^31^). The U2AF and SF1 splicing factors bind the 3' splice site in the H- and E-complexes and subsequently stabilize addition of SF3B1 to form the A-complex. With increasing amounts of NSC 194308, the C- complexes are lost and B complexes are drastically reduced. The A-like complexes still form, but do not accumulate to the same levels as induced by an SF3B1-directed inhibitor of the spliceosome (spliceostatin-A)^32,33^. More pre-mRNA migrates in the region of the gel associated with E-complexes, which is the point that U2AF joins the intron, along with the H-complexes that form in a splicing-independent manner. Because E- and A-complexes precede U2AF dissociation^34,35^, we concluded that NSC 194308 inhibits the pre-mRNA splicing process by stalling spliceosome assembly at a U2AF-dependent checkpoint prior to tri-snRNP recruitment and catalytic activation.

### NSC 194308 targets an inter-RRM cleft of the U2AF2 subunit

Because we identified NSC 194308 in a screen using the ternary U2AF1^S34F^–U2AF2–SF1–*DEK(-3U)* ^Fl^RNA ribonucleoprotein, the exact target of the compound was uncertain. As a step towards identifying the binding site of NSC 194308, we used CANDOCK, a fragment-based docking software that leverages a knowledge-based forcefield for pose scoring^36^. Although a structure of the intact, three protein complex was unavailable, we compared the scores of NSC 194308 docking on the piecewise structures of the human U2AF1, SF1, and two conformations of the U2AF2 subunit, an open RRM1/RRM2 optimally positioned to recognize uridine-rich RNA^37,38^ or a closed RRM1/RRM2 arrangement that predominates in the absence of RNA^38^ (**Fig. 3a, b, Supplementary Table 2**). The most favorable score for binding the hit enhancer was obtained for the open U2AF2 compared to closed U2AF2 or other components of the complex. The top ranked binding site wedged the NSC 194308 molecule between the tandem RRMs of the open U2AF2 conformation and near the bound oligonucleotide, consistent with a structural mechanism of action that stabilizes this major RNA-associated conformation.

We tested the putative binding site of NSC 194308 between the U2AF2 RRMs by monitoring surface plasmon resonance using a BIAcore T200 (**Fig. 3**). The U2AF2 RRM1/RRM2 regions (U2AF2^1,2L^, residues 141-343) was tethered to a sensor chip slide *via* an N-terminal GST tag and serial dilutions of NSC 194308 injected over the surface. The RNA site was omitted due to its rapid dissociation and large response compared to the compound. Although the rapid rates of compound binding the GST-U2AF2^1,2L^ protein approached the reliable limit of the BIAcore instrument, the steady-state responses of the concentration series could be fit in replicate to obtain an average apparent equilibrium dissociation constant (K_D_) of approximately 150 μM (**Fig. 3c-d**), slightly weaker than the estimated EC_50_ for the ribonucleoprotein complex consistent with the absence of the RNA site. The steady-state responses of NSC 194308 injections over individual, GST-tagged RRM1 or RRM2 domains were less than observed for the intact RRM1/RRM2 region and failed to saturate at the highest concentration (estimated K_D_ >1 M) (**Fig. 3d**). In agreement with the docking result, NSC 194308 bound the U2AF2 subunit in a manner that depended on the integrity of the tandem RRMs.

We further probed the location of the NSC 194308 binding site by comparing the EC_50_s of structure-guided U2AF2 mutants in FP assays with the *DEK(-3U)* ^Fl^RNA, which enabled inclusion of the RNA ligand. The potency of the hit compound for U2AF2^1,2L^ – *DEK(-3U)* ^Fl^RNA was similar as for the ternary U2AF1^S34F^–U2AF2–SF1–*DEK(-3U)* ^Fl^RNA complex (**Fig. 3e**). This result supported the conclusion that NSC 194308 acts by binding the U2AF2 subunit. We then compared an internal deletion of a linker connecting the U2AF2 RRM1 and RRM2 (residues 258-257, **Fig. 3f**)^37–39^. The linker deletion reduced the sensitivity of the U2AF2 variant to NSC 194308 (EC_50_ 290 μM). We next prepared a double mutation (D215R/G319R) that is known to promote the closed inter-RRM conformation of U2AF2^38^. The D215R/G319R mutation nearly abolished the sensitivity of the U2AF2 variant to NSC 194308 (**Fig. 3g**). Thirdly, we introduced an electronegative G154E mutation that is expected to repel NSC 194308 from the docked site of U2AF2 (**Fig. 3h**). We found that the G154E mutation reduced the response of U2AF2 to NSC 194308, in agreement with the binding site of the docked model. Altogether, we concluded that NSC 194308 binds near G154 at the major U2AF2 RRM1/RRM2 interface with uridine-rich splice site RNA.

### Charged and hydrophobic groups of NSC 194308 contribute to U2AF2-targeted activity

With the knowledge that NSC 194308 targets the U2AF2 subunit, we investigated the importance of the distinct chemical groups of the compound for inhibition of pre-mRNA splicing in nuclear extract (**Fig. 4**, **Supplementary Fig. 3**). We first synthesized a variant that omitted the anionic thiosulfate (*(-)*-*cis*-myrtanyl acetamidine) (Online Methods, **Supplementary Fig. 3a-b**). Removal of this functional group nearly abolished the anti-splicing activity of the compound. Next, we compared variations in the hydrophobic group by testing different compounds obtained from the NCI repository (**Fig. 4a**). Reducing the hydrophobic portion of the compound to three carbons (NSC 187511) or to the minimal hexane scaffold (NSC 194285) significantly reduced the activity of the compound, whereas altering the shape by replacement with a globular adamantane (NSC 187514) had little effect (**Fig. 4, Supplementary Fig. 3e**). Conversely, either extending the hexane group by a four-carbon linker from the charged group or adding additional hydrophobic methyl groups restored similar potency as the NSC 194308 parent. We concluded that both the anionic thiosulfate and hydrophobic contacts contributed to the anti-splicing activity of the NSC 194308 hit compound.

**Fig. 4.**
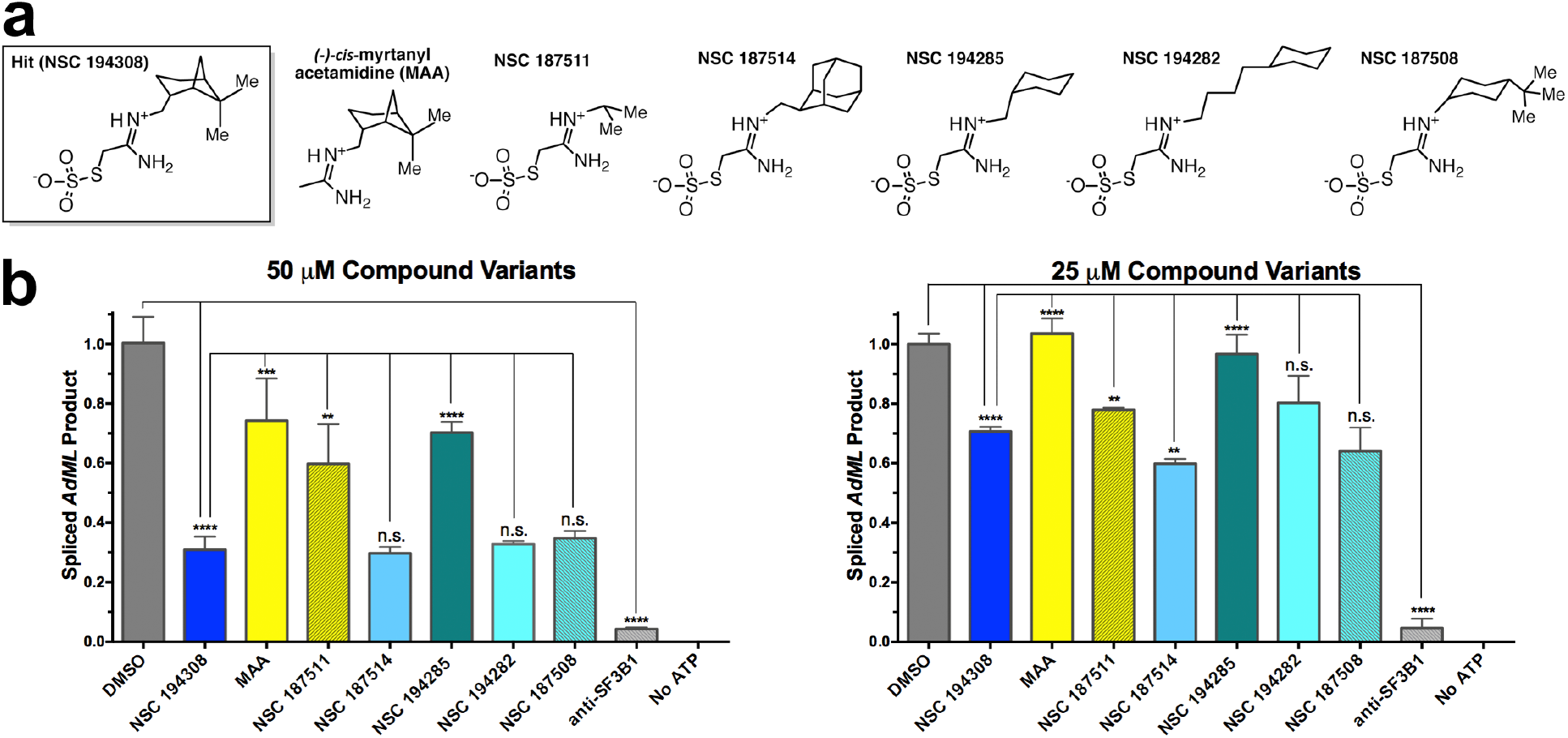
Comparison of NSC 194308 variants shows that anti-splicing activity depends on the thiosulfate and is sensitive to variations of the hydrophobic group. (**a**) Chemical structures of hit compound variants. The (-)-*cis*-myrtanyl acetamidine was synthesized *de novo* (Compound 3, Star Methods and Supplementary Fig. 2) and other compounds were obtained from the NCI Developmental Therapeutics Program. Me, Methyl. (**b**) Comparison of the *in vitro* splicing activities by qRT-PCR of the spliced *AdML* product from HeLa nuclear extract treated with either 50 μM (left) or 25 μM (right) compound variants. The products were normalized to a mock-treated negative control (1% v/v DMSO) and compared to a pladienolide B (anti-SF3B1) positive control. The average values of three replicates are plotted. Unpaired, two-tailed t-tests with Welch’s correction: *, *p* < 0.05; **, *p* < 0.005; ***, *p* < 0.0005; ****, *p* < 0.00005.

### NSC 194308 blocks U2AF2-sensitive splicing in cells

With the knowledge that NSC 194308 targets the U2AF2 subunit, we asked whether treatment with the hit compound would modulate U2AF2-dependent splicing in cells (**Fig. 5**). We treated HEK 293T cells with increasing concentrations of NSC 194308 and isolated RNA after five hours of treatment, a timepoint at which nearly all cells remained viable. To compare the effects of reduced U2AF2 levels, we transfected HEK 293T cells with U2AF2-targeted siRNA and isolated RNAs either two or three days following transfection, which provided sufficient time for U2AF2 levels to decrease. First, we confirmed that U2AF2 levels were reduced by siRNA and remained constant after NSC 194308 treatment under the assay conditions (**Supplementary Fig. 4**). Then, we used qRT-PCR to analyze the NSC 194308-associated changes in the levels of three well-characterized transcripts (*CCND1*, *DUSP11,* and *HTATSF1*) that are subject to regulation by SF3B1 inhibitors and U2AF2-family proteins^12,33,40^ (**Fig. 5a-c**). The results shown in **Fig. 5** represent multiple biological replicates and are averages of three technical replicates normalized to *GAPDH*. The relative abundance of *CCND1*, *DUSP11,* and *HTATSF1* was reduced by treatment with NSC 194308 to a similar extent as by knockdown of *U2AF2*. The levels of *CCND1* were most sensitive to the compound (IC_50_ ~10 μM) compared to the other transcripts (IC_50_ ~75 μM). These apparent potencies of the compound for modulating transcript levels in cells agreed with the values for inhibiting pre-mRNA splicing *in vitro.*

**Fig. 5.**
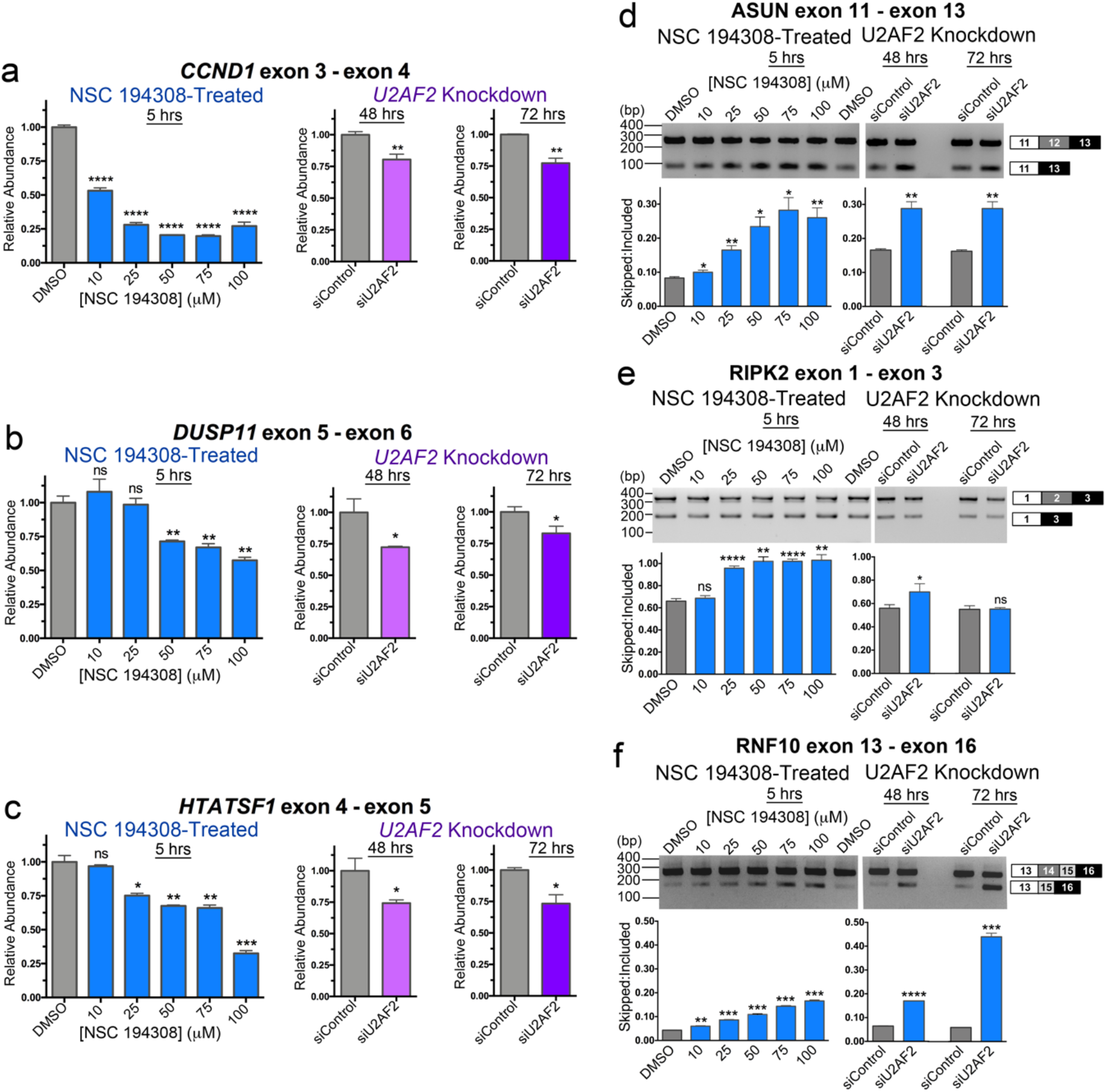
NSC 194308 alters splicing of U2AF2-sensitive transcripts in HEK 293T cells. (**a** - **c**) qRT-PCR of gene transcripts with known responsiveness to U2AF2 family members. NSC 194308-treated samples (blue, left) are compared with samples treated with *U2AF2*-targeted Stealth™ siRNAs (or control siRNA) for either two days (magenta, center) or three days (purple, right). The products of the indicated gene transcripts were normalized to GAPDH controls. For Figures 5 and 6, the qRT-PCR results are the averages of three technical replicates. (**d** - **f**) RT-PCR of the indicated U2AF2-regulated gene transcripts following either treatment with NSC 194308 (left) or *U2AF2* knockdown (right). A mocked-treated control (0.1% v/v DMSO) matches the NSC194308 solvent. The average exon-skipped:included ratios and standard deviations of background-corrected bands from three reactions (six for DMSO control) are plotted below representative ethidium-bromide-stained agarose gels. The expected PCR products are schematized to the right. Immunoblots of samples used for RNA isolation are shown in Supplementary Fig. 6. No products were detected in negative control reactions. All results represent multiple biological replicates. Unpaired, two-tailed t-tests with Welch’s correction: *, *p* < 0.05; **, *p* < 0.005; ***, *p* < 0.0005; ****, *p* < 0.00005.

Although detected at the majority of 3' splice sites^41,42^, U2AF2 extensively regulates alterative splicing (e.g.^42–44^). To investigate whether NSC 194308 specifically modulated U2AF2-dependent alternative splice sites, we next used reverse transcription (RT)-PCR to examine the effects of the compound on three representative *U2AF2*-dependent exon-skipping events (**Fig. 5d-f**). Following knockdown of *U2AF2* in HEK 293T cells, we noted robust increases in the skipping of *ASUN* (also called *INTS13*) and *RNF10* exons, whereas *RIPK2* changes were subtle, in agreement with previous observations using HeLa cells^42^. Remarkably, addition of NSC 194308 increased skipping of the *U2AF2*-sensitive exons in the spliced products. The similar effects of *U2AF2* knockdown and NSC 194308 treatment were consistent with the ability of the compound to stall *in vitro* spliceosome assembly at the stages involving U2AF2 (**Fig. 2**).

### Cells expressing mutant U2AF1 have increased sensitivity to NSC 194308

Considering the premise that human cells expressing mutant splicing factors are sensitized to splicing modulators, we asked whether NSC 194308 treatment would selectively decrease the survival of K562 leukemia cells expressing the MDS-associated S34F mutant of U2AF1. First, we tested the NSC 194308-sensitivity of a cell line with a stably integrated doxycycline-inducible, FLAG-tagged S34F-mutant *U2AF1* compared to a FLAG-tagged wild-type *U2AF1* control for overexpression (**Supplementary Fig. 5a**). Treatment with NSC 194308 significantly reduced the survival and lowered the IC_50_ (*p*-value 0.0047 in unpaired t-test with Welch’s correction) of S34F mutant-induced K562 cells compared to the wild-type induced control cells (**Fig. 6a**). Notably, the apparent IC_50_ of the K562 cells overexpressing the MDS-associated S34F mutation occurred at much lower NSC 194308 concentrations than the purified ribonucleoprotein complex (approximately 5 μM compared to 100 μM), suggesting that slight perturbations of the essential U2AF2 protein could have a larger functional impact in the context of prior insults to the pre-mRNA splicing pathway. Next, to reflect the levels of untagged U2AF1 expression expected in cancer cells, we edited either the S34F *U2AF1* mutation or a wild-type control into a K562 cell line using CRISPR methods. Successful introduction of the mutation was verified by genomic sequencing and supported by S34F-associated changes in splicing (**Supplementary Fig. 5b-d**). Following NSC 194308 treatment, the survival and IC_50_ of the edited cell line expressing physiological levels of S34F-mutant U2AF1 also were significantly reduced (*p*-value 0.0065) compared to the wild-type counterpart (**Fig. 6a**). The selective killing of S34F U2AF1-expressing cells by NSC 194308 was similar in magnitude to previously observed effects on analogous cell lines of an SF3B1 inhibitor, sudemycin D1 (approximately 2-4-fold selection)^10^.

**Fig. 6.**
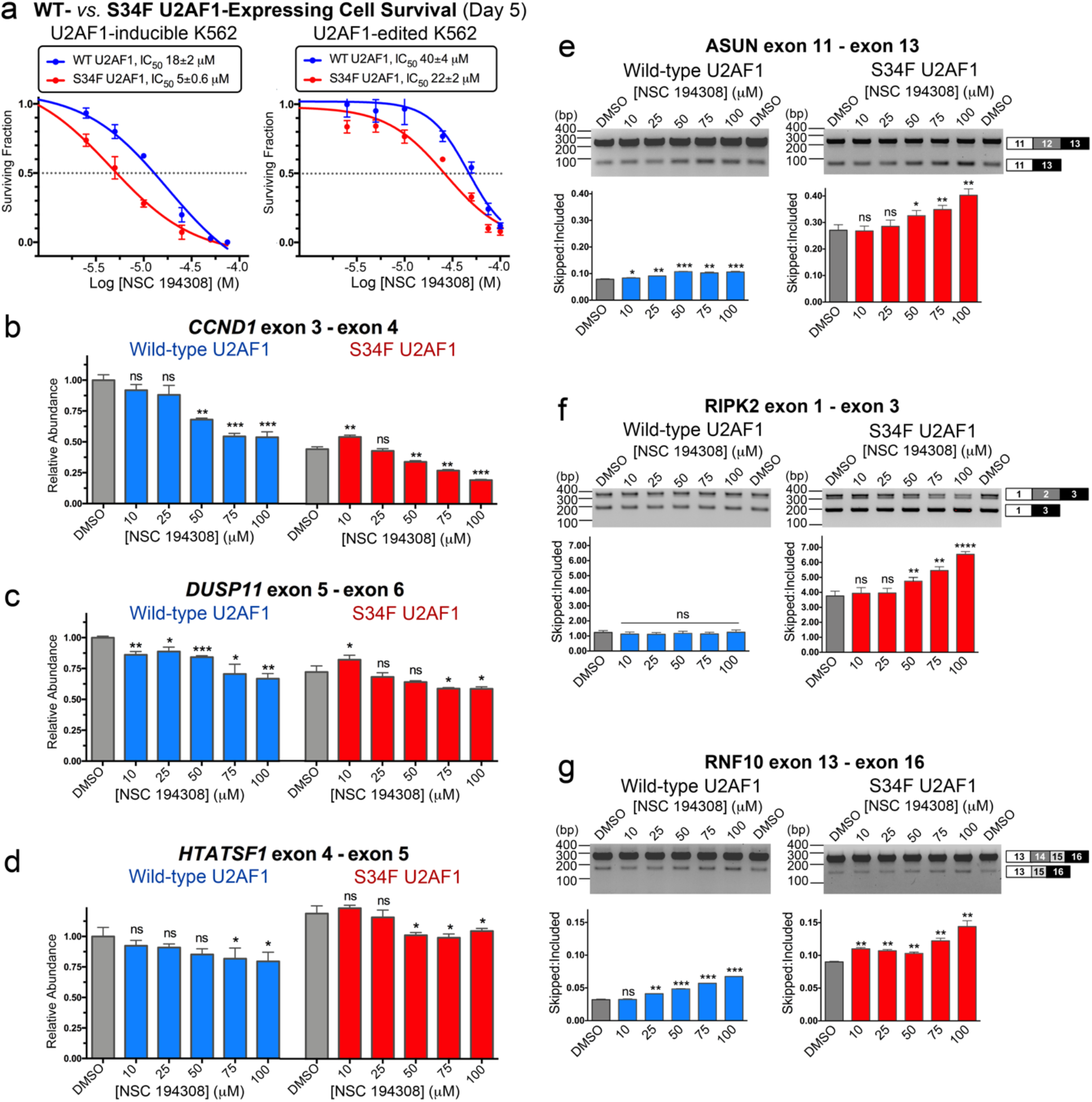
NSC 194308 preferentially kills K562 leukemia cells expressing MDS-associated S34F-mutant U2AF1 and alters splicing in these cells. (**a**) Fraction of viable cells (detected by trypan blue staining) normalized to a mock-treated sample (0.1% v/v DMSO) and plotted *versus* the log_10_ concentration of NSC 194308. *Left:* K562 cells with stably integrated, doxycycline-inducible WT or S34F-mutant *U2AF1* cultured with increasing concentrations of NSC 194308 concurrently with doxycycline (250 ng mL^−1^) for five days following two days of initial induction. *Right:* K562 cells with endogenous *U2AF1* gene edited by CRISPR technique. The WT or S34F mutant cells were treated with the indicated concentrations of NSC 194308 for five days. The inhibitory concentrations (IC_50_) at half-maximal cell survival are inset above. (**b** - **d**) qRT-PCR (normalized to GAPDH control of the WT sample) or (**e** - **g**) RT-PCR of the indicated gene transcripts from *U2AF1*-edited cells treated for seven hours with NSC 194308. The average exon-skipped:included ratios and standard deviations of background-corrected bands from three reactions (six for DMSO control) are plotted below representative ethidium-bromide-stained agarose gels. Immunoblots of samples used for RNA isolation are shown in Supplementary Fig. 7. Wild-type (WT), blue; S34F-mutant, red. Unpaired, two-tailed t-tests with Welch’s correction compare the indicated NSC 194308-treated sample with the matching WT or S34F mutant DMSO control: *, *p* < 0.05; **, *p* < 0.005; ***, *p* < 0.0005; ****, *p* < 0.00005.

With the knowledge that NSC 194308 inhibits U2AF2-dependent splicing and that U2AF1 is a heterodimeric partner of U2AF2, we thought that dysregulated splicing was likely to contribute to the sensitivity of S34F U2AF1-expressing cells to the compound. To explore this possibility, we examined representative transcripts isolated from edited K562 cell lines following seven hour treatments with increasing concentrations of NSC 194308 (**Fig. 6**). In general, the changes in transcripts from the NSC 194308-treated K562 leukemia cells were similar to those of RNAs from HEK 293T cells treated with NSC 194308 or *U2AF2*-targeted siRNA shown in **Fig. 5** and described in reference^42^. In some cases, the presence of the S34F mutation of U2AF1 appeared to exacerbate or alter the NSC 194308-induced changes in gene expression. In particular, treatment with the compound further decreased the levels of *CCND1* transcript or the exon-included *RIPK2* splice form, which were already expressed at lower levels in the S34F-mutated cells than in the wild-type counterpart. We concluded that aggravation of splicing defects by NSC 194308 is a likely mechanism of action for its preferential killing of cells expressing mutant U2AF1.

## Discussion

In this study, we demonstrated that the U2AF2 splicing factor and its RNA binding properties can be targeted *de novo* by a small molecule. The success of our small-scale classical screen implies that expansion of such an approach would identify additional U2AF2 modulators. We identified the binding site of NSC 194308 between the U2AF2 RRMs by a combination of docking, mutagenesis, and binding assays. This result is significant because the current repertoire of splicing factors targeted by small molecules is limited. Despite a plethora of recurrent mutations in the early components of the pre-mRNA splicing machinery across multiple cancer types, the major class of clinically advanced spliceosome inhibitors universally target the same site of the splicing factor SF3B1 (reviewed in ^13^). Although inhibitors of pre-mRNA splicing have been identified from chemical library screens of pre-mRNA splicing in nuclear extracts or cells, often the potencies are low and identifying the target for optimization is challenging due to the complexities of >100 participants in the splicing reaction. Antisense oligonucleotides show promise as modulators of specific splice sites, yet the ambiguity of the exact cancer drivers coupled with the difficulty of oligonucleotide delivery are on-going challenges confronting clinical applications. An innovative small molecule directed at U2AF2 offers a tool to dissect roles of this key splicing factor in normal and dysfunctional gene expression. In broader terms, optimized U2AF2 inhibitors have the potential to exploit the widespread vulnerability of dysregulated pre-mRNA splicing in a cancers, by analogy with the successful killing of spliceosome-defective cells by SF3B1 inhibitors, regardless of the presence of SF3B1 mutations^5–8,10,12^. Already, we have demonstrated that U2AF2 modulators preferentially kill a leukemia cell line carrying the S34F mutation of *U2AF1* that is commonly found in MDS, which currently lacks a chemotherapeutic cure.

An equally significant outcome of this project is the power of stabilizing stepwise checkpoints of spliceosome activation as a means to alter the splicing process. Notably, all three families of SF3B1-targeted splicing inhibitors stall spliceosome assembly at the A-stage near the time of U2AF2 release^32,33,45,46^, although such inhibitors also can interfere with later stages of splicing^47^. Similarly, an inhibitor of *in vitro* splicing recently identified in a chemical library screen was found to block the conversion of B-stage spliceosomes to the activated B^act^ configuration^48^. Conversely, RNA-targeted enhancers of U1 small nuclear ribonucleoprotein (snRNP) association have been leveraged to promote splicing of the defective survival of motor neuron copy for treatment of spinal muscular atrophy ^49,50^, and by analogy, our findings raise the question of whether splicing would be stalled at elevated concentrations of these compounds. Here, our discovery of NSC 194308 represents the first small molecule to arrest spliceosomes at the U2AF2-dependent E- to A-complex transition as a potential tool for studies of pre-mRNA splicing mechanisms. Identifying the binding site for NSC 194308 on U2AF2, together with establishing its general structure-activity relationships, lays a guiding foundation for future optimization of the compound’s potency in drug-like applications.

The vast majority of functional pathways are tightly regulated by critical checkpoints. This work illustrates the feasibility of blocking specific pathways by using small molecules to stabilize checkpoints of macromolecular assemblies, highlighting a potential broad strategy for the development of new molecular tools and therapeutics.

## Acknowledgements

We are grateful to Prof. A. V. Smrcka (U. Michigan) for advice designing the screen. This work was supported by grants from the National Institutes of Health (NIH R01 GM070503 to C.L.K. and R01 GM122279 to M.S.J.), the National Science Foundation (NSF CHE-1900050 to A.F.), UR Ventures (to C.L.K.), and the Edward P. Evans Foundation (to C.L.K., M.J.W., and T.A.G.). Work by R.S. was supported by an NIH Director**’** s Pioneer Award (1DP1OD006779), Clinical and Translational Sciences (NCATS) Award (UL1TR001412), and NCATS ASPIRE Design Challenge Award. A.M. was supported by NIH training grant T32 GM08646. Z.F. was supported by the National Library of Medicine (T15 LM012495). An NSF Major Research Instrumentation grant (CHE-1725028) supported the X-ray diffractometer.

## Author Contributions

C.L.K., R.C. and M.J.P. conceived experiments in collaboration with all authors. R.C. screened small molecule libraries. C.F.F. accomplished *in vitro* experiments with guidance from J.L.J., M.J.P., and C.L.K. A.J.M. and M.S.J. analyzed spliceosome assemblies. Z.F. and R.S. performed computational docking. G.A., W.W.B. and A.J.F. carried out synthesis, purification, and characterization of compounds. S.R., M.J.W. and T.A.G. constructed cell lines. M.J.P. accomplished biological experiments. C.L.K. wrote the paper with contributions from R.C. M.S.J., G.A., A.J.F., M.J.W., T.A.G. and input from all authors.

## Additional Information

Supplementary information is available for this paper.

## Methods

### Preparation of proteins and RNAs for binding assays

The ternary protein complex of human U2AF1^S34F^ (wild-type or S34F mutant, residues 1-193 of NCBI RefSeq NP_006749), U2AF2 (residues 85-471 of NP_001012496), and SF1 (residues 1-255 of NP_004621) was prepared as described^29^. In brief, the U2AF1 and U2AF2 subunits were co-expressed in *Escherichia coli* and the tags were removed. The SF1 subunit was mixed with the heterodimer and the ternary complex was isolated by size exclusion chromatography in a buffer containing 25 mM HEPES pH 6.8, 150 mM NaCl, 3% glycerol, 20 μM ZnCl_2_, 3 mM βME and 0.5 mM TCEP. The RNA recognition motif region of the U2AF2 subunit (residues 141-342) was prepared as described^37^. The combined yield of protein prepared from approximately 12 liters of *E. coli* host cells (6 L co-expressing the human heterodimer plus 6 L expressing SF1) was used for each screen (for inhibitors or enhancers).

The RNA oligonucleotides were synthesized and HPLC purified by Dharmacon™ (Horizon Discovery). The *DEK(-3U/-3C)* RNA oligonucleotide sequences are 5’-UAAGAAAUACUAAAUUAAUUUC**(U/C)** AG AAAAGAGUCU. Where indicated, a fluorescein (Fl) was tethered *via* a 6-methylene carbon linker to the 5' terminus of the oligonucleotide.

### Fluorescence polarization (FP) screens for U2AF–RNA modulators

Data of the screens are summarized in **Supplementary Table 1**. Positive and negative control samples were plated in 24 replicates in 384-well flat bottom clear polystyrene plates (Corning, USA). A concentration of protein (80 nM) was chosen preceding the exponential phase of the binding curve of the low affinity ^Fl^*DEK(-3U)* RNA site (20 nM). Excess U2AF1^S34F^–U2AF2–SF1 protein (1.3 μM) served as the positive control for enrichment of the complex and matching solvent (1% v/v DMSO) as the negative control. The samples were incubated at room temperature for 60 min, an empirically determined timepoint to equilibrate the reading. The FP was measured at 520 nm using an EnVision plate reader following excitation at 490 nm.

The NCI Diversity Set V library (1,593 compounds) in 96-well format (10 mM stock in DMSO) was transferred to 384-well plates and diluted to 1 mM in DMSO using a JANUS Varispan Automated Workstation (PerkinElmer). The U2AF1^S34F^–U2AF2–SF1 protein and ^Fl^*DEK* variant oligonucleotide were plated at a total 20 μL reaction volume per well. The JANUS pintool was used to transfer 100 nL aliquots from the library plate to the sample wells and achieve a 5 μM final concentration of each compound per well. The unlabeled RNA, excess protein, or DMSO were added to the appropriate positive and negative control wells as described above. The sample plates were incubated at room temperature for 60 min, the timepoint at which the Z’-factor had plateaued. Then, FP was measured using the EnVision plate reader. The quality of the HTS was assessed using the statistical Z-score^30^, for which μ_s_ and μ_C-_ are the respective means of the samples and negative control signals and σ_s_ and σ_C_- are the standard deviations:

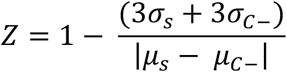

The compounds that increased the FP of the sample wells by >75% compared to the signal of the positive controls were considered “hits”. To test the reproducibility of each hit compound’s effect compared to appropriate positive and negative controls, these initial hits were cherry-picked using the JANUS workstation and the FP measurements were replicated in triplicate. The FP dose-response curves of reproducible U2AF1^S34F^–U2AF2–SF1–^Fl^*DEK* RNA modulators were measured using a Fluoromax-3 fluorimeter. The compound was titrated into a mixture of protein and RNA in the cuvette. The DMSO solvent was maintained at <2% of the total sample volume. The sample was excited at 490 nm and the anisotropy measured at 520 nm. The data were fit with a sigmoidal dose response model using Prism (GraphPad Software).

### Synthesis of hit compound NSC 194308 and variant

All reactions were carried out under an argon atmosphere with magnetic stirring, unless otherwise noted. Reagents were used as obtained from commercial suppliers without further purification. Dry solvents were dried for at least 24 hours over activated 3 Å molecular sieves before use in any reactions. Reactions were monitored by thin layer chromatography (TLC) separation on pre-coated silica gel 60 F254 glass-supported plates (Millipore-Sigma). The TLC plates were visualized under UV light then stained using *p*-anisaldehyde/sulfuric acid solution and gentle heating. NSC 194308 was dissolved in DMSO solvent at 10 mM stock concentration, stored at −80° C, and serial diluted in DMSO or matching buffers as needed immediately prior to each experiment. The ^1^H NMR spectra were recorded at room temperature on a 500 MHz Bruker Avance spectrometer. Chemical shifts are given in parts per million (ppm) referenced to solvent residual proton resonance (δ = 3.31 ppm for d4-MeOD or δ = 2.50 ppm for d6-DMSO). NMR data are reported as: chemical shift, multiplicity (s = singlet, d = doublet, t = triplet, q = quartet, m = multiplet, dd = doublet of doublets, dq = doublet of quartets, br = broad), coupling constants (*J*) given in Hz, and integration. The ^13^C NMR spectra were recorded at room temperature on a 125 MHz Bruker Avance spectrometer, with proton decoupling. Chemical shifts are given in parts per million (ppm) from referenced to solvent carbon resonance (δ = 49.00 for d4-MeOD).

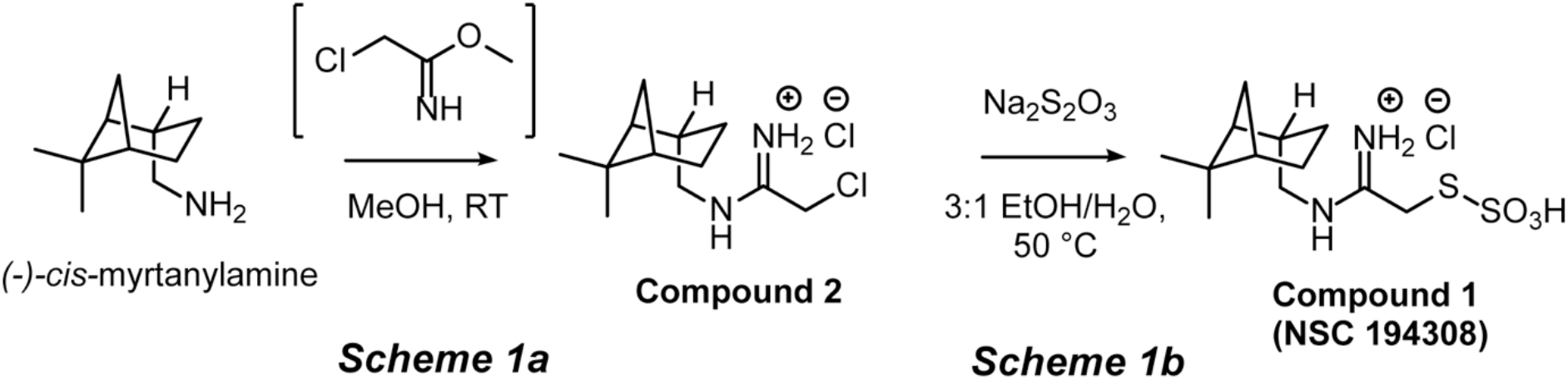

#### Synthesis of Compound 2 from (-)-cis-myrtanylamine (Scheme 1a)

Metallic Na (23 mg, 1 mmol) was added to a flame-dried round bottom flask containing dry MeOH (5 ml), and an exothermic reaction took place with evolution of H_2_ gas. This solution was stirred for 1 h at room temperature, upon which chloroacetonitrile (226 mg, 2.5 mmol) was added in a drop-wise fashion. This new solution was allowed to react at room temperature for 2 hours. Then, *(-)*-*cis*-myrtanylamine (306 mg, 1 mmol) was added dropwise, and the pH adjusted to roughly 1 with 3.3 M ethanolic HCl (circa 1 ml). The solution was allowed to react for 16 h at room temperature. The reaction mixture was concentrated, and isopropyl alcohol (iPrOH) (30 ml) was added, and the solids were triturated and filtered. The filtrate was concentrated to roughly half the initial volume (approximately 15 ml) and diethyl ether (15 ml) was added and the mixture was left to stand for 15 min. The product **Compound 2** (2-chloro-*N-(-)-cis*-myrtanyl acetamidinium hydrochloride) was filtered off as a pink powder, and dried under vacuum (371 mg, 70% yield). The crystal structure of Compound 2 was determined as described below and is shown in **Supplementary Fig. 2a**. **^1^H NMR (500 MHz, d6-DMSO)** δ 10.07 (br s, 1H), 9.54 (br s, 1H), 9.12 (br s, 1H), 4.41 (s, 2H), 3.22 (t, J = 6.9 Hz, 1H), 2.35 – 2.31 (m, 2H), 1.95 – 1.89 (m, 5H), 1.46 – 1.37 (m, 1H), 1.18 (s, 3H), 1.01 (s, 3H), 0.87 (d, *J* = 9.4 Hz, 1H) ppm.

#### Synthesis of Compound 1 from Compound 2 (Scheme 1b)

To a stirred solution of **Compound 2** (66 mg, 0.25 mmol) in ethyl (Et) alcohol (0.75 ml) in an Erlenmeyer flask was added a solution of Na_2_S_2_O_3_ (40 mg, 0.25 mmol) in H_2_O (0.25 ml). The combined solution was heated to 50 ^o^C for 1 h. Upon full consumption of the starting material by TLC monitoring, the solution was allowed to cool to room temperature. The solution was kept acidic by addition of 1 drop of concentrated hydrochloric acid (37%) while stirring, and afterwards the mixture was allowed to stand at room temperature for 15 min without stirring. The product **Compound 1** (NSC 194308) was filtered off as a white powder, and dried under vacuum (57 mg, 67% yield). **^1^H NMR (500 MHz, MeOD)** δ 9.25 (br s, 1H), 8.94 (br s, 1H), 8.71 (br s, 1H), 3.98 (s, 2H), 3.30 – 3.21 (m, 2H), 2.61 – 2.31 (m, 2H), 2.22 – 1.82 (m, 5H), 1.54 (dt, J = 20.5, 8.5 Hz, 1H), 1.24 (s, 3H), 1.07 (s, 3H), 0.98 (d, J = 9.8 Hz, 1H) ppm. **^13^C NMR (125 MHz, d4-MeOD)** δ 167.5, 49.8, 44.9, 42.5, 40.5, 39.6, 35.8, 33.9, 28.3, 26.9, 23.5, 20.7 ppm (**Supplementary Fig. 2b**).

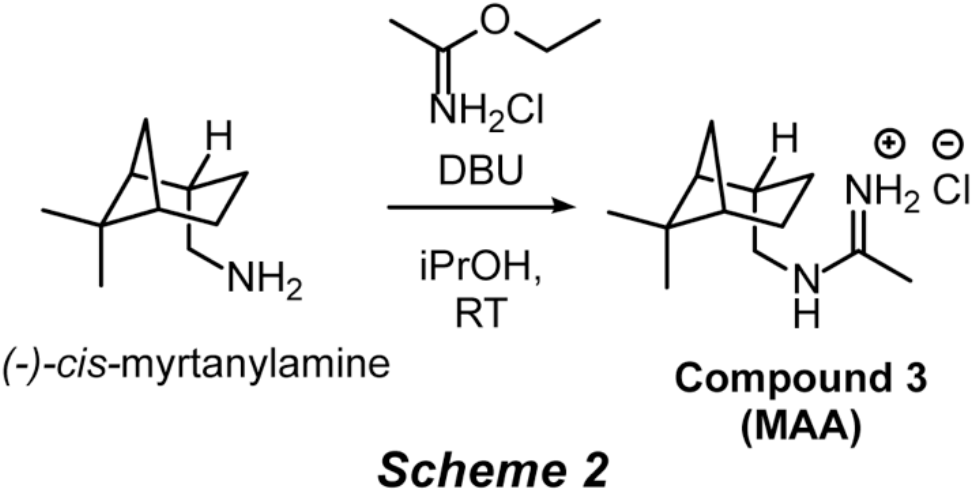

#### Synthesis of Compound 3 from (-)-cis-myrtanylamine (Scheme 2)

1,8-Diazabicyclo[5.4.0]undec-7-ene (304 mg, 2 mmol) was added to a stirred solution containing dry iPrOH (2 ml), ethyl acetimidate hydrochloride salt (248 mg, 2 mmol) and *(-)-cis*-myrtanylamine (153 mg, 1 mmol) and the solution was allowed to stir at room temperature for 20 h. Water (20 ml) was added along with Et_2_O (20 ml) and the reaction was transferred to a separating funnel. The aqueous layer was extracted twice more with Et_2_O (20 ml each) and the combined organic layers were dried over MgSO_4_ and concentrated. The crude oil was dissolved in iPrOH (20 ml) and acidified with 3.3 M HCl in EtOH until a pH of roughly 1 was obtained. The turbid mixture was concentrated and the solid imidinium salt was triturated with EtOAc (20 ml) and filtered to yield white flakes of **Compound 3** (*N-(-)-cis*-myrtanyl acetamidinium hydrochloride, MAA). The compound was then recrystallized from 19:1 acetone/iPrOH to give lustrous white needles (109 mg, 51% yield). The crystal structure of Compound 3 was determined as described below and is shown in **Supplementary Fig. 3a**. **^1^H NMR (500 MHz, d4-MeOD)** δ 3.28 – 3.16 (m, 2H), 2.46 (dtd, J = 11.8, 6.1, 1.8 Hz, 1H), 2.42 – 2.32 (m, 1H), 2.22 (s, 3H), 2.11 – 1.90 (m, 5H), 1.59 – 1.43 (m, 1H), 1.24 (s, 3H), 1.07 (s, 3H), 0.98 (d, J = 9.8 Hz, 1H) ppm. **^13^C NMR (125 MHz, d4-MeOD)** δ 166.1, 49.9, 44.9, 42.5, 40.5, 39.6, 34.0, 28.3, 26.9, 23.5, 20.7, 18.8 ppm (**Supplementary Fig. 3b**).

#### Crystal Structure Determinations of Compound 2 and Compound 3

Compounds 2 and 3 were each crystallized from 19:1 acetone/iPrOH. Data sets were collected at 100 K using a Rigaku XtaLAB Synergy-S Dualflex diffractometer equipped with a HyPix-6000HE HPC area detector and PhotonJet (Cu) X-ray source. Structures were determined using ShelXT^51^, refined using ShelXL^52^, and converged to R_factors_ of 0.118 and 0.039 for *I*>2σ(*I*). Structures were manipulated and figures generated using Olex2^53^ (**Supplementary Figs. 2a and 4a**).

### In vitro splicing assays using nuclear extract

The DNA templates for *AdML(−3C)* and *AdML(-3U)* substrate pre-mRNAs have been described^54,55^. For RT-qPCR splicing analysis, the pre-mRNAs were prepared using the mMESSAGE mMACHINE™ T7 transcription kit (Thermo Fisher Sci.) and purified using the MEGAclear™ transcription clean-up kit (Thermo Fisher Sci.). The *in vitro* splicing reaction used HeLa nuclear extract (ProteinOne) with a similar protocol as described^56^, except that the reactions were incubated at 30 °C for one hour, heat-inactivated at 60 °C for 10 min, diluted two-fold, and finally stored at −80 °C. The spliced products were diluted to 1/40^th^ of the total PCR reaction volume and quantified by Taqman qRT-PCR as described^56^.

For denaturing gel splicing analysis, G(5′)ppp(5′)G-capped and ^32^P-UTP body-labeled pre-mRNA was created using standard T7 run-off transcription and gel purified. Nuclear extract was prepared from HeLa cells cultured in MEM/F12 1:1 and 5% (v/v) newborn calf serum and stored in 20 mM Tris pH 7.9, 0.1 M KCl, 0.2 mM EDTA, 20% (v/v) glycerol, 0.5 mM DTT^57^. The *in vitro* splicing reactions contained 10 nM pre-mRNA, 60 mM potassium glutamate, 2 mM magnesium acetate, 2 mM ATP, 5 mM creatine phosphate, 0.05 mg/mL tRNA, 40% (v/v) HeLa nuclear extract and increasing amounts of drug solved in 1% (v/v) DMSO, and were incubated at at 30°C for 60 min. Following phenol:chloroform extraction and ethanol precipitation, RNAs were separated by electrophoresis in a 15% (v/v) polyacrylamide denaturing gel with 7 M urea in 1X TBE (45 mM Tris-borate, 1 mM EDTA). Gels were run at 35 W for 2 hours and visualized by phosphorimaging with a Typhoon Scanner (Molecular Dynamics).

For native gel analysis of spliceosome assembly, aliquots of the same splicing reactions were mixed with an equal volume of native loading buffer (20 mM Trizma base, 20 mM glycine, 25% (v/v) glycerol, 0.05% (w/v) cyan blue, 0.05% (w/v) bromophenol blue, 1 mg/mL heparin sulfate. Splicing complexes were separated by electrophoresis in a 2.1% (w/v) low-melt agarose native gel in 20 mM Tris, 20 mM glycine. Gels were run at 72 V for 3 h 50 min, vacuum-dried onto Whatman paper for 45 min at 65 °C, and visualized by phosphorimaging as described above.

### Parameterization of docking software and preparation of structures

NSC194308 was docked to the separate structures of the U2AF1, U2AF2, and SF1 components using CANDOCK, a fragment-based docking protocol that implements knowledge-based statistical functions for pose minimization and scoring^36^. Unique chains of interest were extracted from the Protein Data Bank (PDB) structure coordinate files in preparation for docking, including PDB ID’s 5EV4^37^, 1JMT^58^, 1K1G^59^, 2YH0^38^, 4FXW^60^, and 4YH8^61^. The Gen3D module of the Open Babel package^62^ was used to generate and optimize the geometry of a 3D model for the NSC 194308 compound. We used COFACTOR to predict candidate ligand binding sites for each protein structure based on homology to templates available in the wwPDB^63,64^. Finally, we used CANDOCK^36^ to assess the fit of the NSC 194308 model coordinates for the candidate binding sites of the protein structures, with otherwise default values. The top minimized poses of docked NSC 194308 for each protein were scored relative to the knowledge-based forcefield as described^36,65^ (**Supplementary Table 2**).

### Biomolecular interaction by surface plasmon resonance

We used a Biacore™ T200 with CM5 sensor chips (GE Healthcare Inc.). We immobilized approximately 11,000 RU of anti-GST in a buffer containing 10 mM HEPES pH 6.8, 150 mM NaCl, 0.2 mM TCEP, 0.005% P20 (HBS buffer). We then immobilized 900 – 1500 RU of GST on the reference surface and a similar amount of each GST-U2AF2 protein on the active surface of the sensor chip. The affinity experiments used matched injection and running buffers of HBS plus 1% (v/v) DMSO that were carefully and reproducibly matched. Each injection of NSC 194308 was repeated in duplicate at a flow rate of 30 μl ^−1^ min and double-referenced to injections of the same buffer. The injection time was 90 s followed by 60 s dissociation. The surface was regenerated by application of 1 M NaCl for 60 s between injections. Data was analyzed using the BIAcore T200 Evaluation software (v3.2, GE Healthcare Inc.). The GST-U2AF2^1,2L^ responses were corrected for a control cell on which GST was immobilized. The responses at equilibrium (R_eq_) were calculated by averaging 5 s of double-referenced data immediately before the end of the injection. The R_eq_ were plotted as a function of the corresponding NSC 194308 concentrations and fit to a one-site binding equation using Prism (GraphPad Software) to obtain the apparent equilibrium dissociation constant.

### Cell culture and transfections

The human embryonic kidney (HEK) 293T cell lines were obtained directly from ATCC^R^ (CRL-3216™). Human embryonic kidney HEK293T cells were cultured in Dulbecco’s modified Eagle media (Gibco™), 10% fetal calf serum (R&D Systems Inc.), 2 mM L-glutamine and penicillin-streptomycin (pen-strep, Gibco™). As needed, cells were split with Trypsin-EDTA (Gibco™). The K562 cell lines expressing doxycycline-inducible 3xFLAG wild-type or S34F U2AF1 have been described^66^. Separately, the S34F mutation or matching wild-type control were introduced into a K562 background using CRISPR methods.

K562 cells from ATCC^R^ (CCL-243™), resuspended in Buffer R (Thermo Fisher Sci.), were gently mixed with the freshly prepared Hifi2-Cas9 (MCLAB) – gRNA (Synthego) ribonucleoprotein complex, single stranded donor oligodeoxynucleotide (ssODN) and electroporated using the Neon Electroporation System (Thermo Fisher Sci.). The clonal lines were expanded by limiting dilution and confirmed to harbor the desired substitution and zygosity by amplicon-based next-generation sequencing. The K562 cell lines were cultured in either RPMI 1640 (Gibco™) for the *U2AF1*- inducible cells or IMDM (Gibco™) for the *U2AF1*-edited cells, supplemented with 10% fetal calf serum, 2 mM L-glutamine and pen-strep. All cells were maintained at 37 °C in a humidified chamber containing 5% CO_2_.

U2AF2 levels were reduced by transfection of HEK 293T cells with Stealth™ siRNA HSS117616 (Thermo Fisher Sci.) and compared to a “lo GC” negative control Stealth™ siRNA using JetPrime^R^ Polyplus-transfection as instructed by the manufacturer. Cells were split after 24 hrs and harvested at two or three days post-transfection.

### Viability Assays of K562 Cells

Inducible K562 cells were plated at 2.5×10^5^ cells mL^−1^ in RPMI 1640 containing 0.25 μg/mL doxycycline (dox) (Takara Bio USA Inc.) in 10 cm plates and incubated at 95% humidity, 5% CO_2_, 37° C. After 48 hrs, cells were diluted in an equal volume of fresh media with dox and plated into 24 well dishes at 1×10^5^ cells mL^−1^. Compound at the indicated concentration, or comparable DMSO control (0.1% v/v), was then added. Live cell numbers were measured after five days by trypan blue exclusion assays as suggested by the manufacturer (Invitrogen). CRISPR-edited K562 cells in IMDM without dox otherwise were compound-treated in an identical manner.

### NSC 194308 treatments for RNA isolation

HEK293T cells were plated in 24 well plates at 1.2×10^5^ cells/mL and incubated overnight. CRISPR-edited K562 were plated in 6 well plates at 2.5×10^5^ cells mL^−1^ prior to treatment with compound. Cells were treated with the indicated amounts of NSC 194308 compound (from 1000x stocks in DMSO) or 0.1% (v/v) DMSO as a negative control. To observe the primary effects of the compound, HEK 293T cells or K562 cells were harvested after five or seven hours of exposure, respectively.

### RT-PCR and qRT-PCR of RNAs from cell lines

Total RNA was isolated from harvested cells and DNase I-treated using the RNeasy Kit (Qiagen). The cDNAs were synthesized using Moloney murine leukemia virus RT with random primers (Invitrogen, Thermo Fisher Sci.). RNA levels of representative transcripts were analyzed in triplicate by quantitative PCR with SYBR™ Green using a Bio-Rad CFX thermal cycler. The RT-qPCR products were quantified by the relative standard curve method and normalized to the level of *GAPDH* RNA. Changes in the alternative splicing were investigated by PCR reactions. The bands were separated by gel electrophoresis on a 2% agarose gel in TBE buffer, visualized by ethidium bromide stain, and imaged using a Bio-Rad Gel Doc XR+. Products of the RT-PCR reactions were quantified using FIJI (ImageJ) densitometry analysis. The primer sequences are provided in **Supplementary Table 3**.

### Immunoblot analysis

Cells were lysed in a buffer containing 50 mM Tris pH 8.0, 10 mM EDTA, 1% SDS, 1 mM DTT, with phosphatase and protease inhibitors. Protein was separated by SDS-PAGE, transferred onto PVDF membranes (Millipore-Sigma), then blocked with 5% milk in TBS-T. Primary antibodies specific for GAPDH (Cell Signaling Technology Inc., Cat. No. 2118), U2AF2 (Millipore-Sigma, Cat. No. U4758), H2B (Cell Signaling Technology Inc., Cat. No. 12364), and U2AF1 (Proteintech^R^ Group Inc., Cat. No. 10334-1-1AP) were diluted in blocking solution at 1:1000 (v/v) ratio. Anti-DYKDDDDK was diluted at 1:2000 (v/v) ratio. Anti-mouse IgG horseradish-peroxidase (Cat. No. NA931) and Anti-rabbit IgG horseradish-peroxidase (Cat. No. NA934), from GE Healthcare Inc., were diluted in blocking buffer at 1:5000 (v/v) ratio. Chemiluminescent signal, using Clarity Western ECL substrate (Bio-Rad), was detected on a Chemidoc™ Touch Imaging System (Bio-Rad).

### Data availability

The coordinates and structure factors are deposited in the Cambridge Crystallographic Data Centre (CCDC) with access codes CCDC 1981586 (Compound 2) and CCDC 1981585 (Compound 3). All other data and materials are available from the authors by reasonable request.

