## Supplementary Data for "A synthetic small molecule stalls pre-mRNA splicing by promoting an early-stage U2AF2–RNA complex"

1) **Supplementary Results:** Screen for hit inhibitors of U2AF – RNA binding.

2) **Supplementary Table 1.** Small molecule screening data.

3) **Supplementary Table 2.** Scores from Candock.

4) **Supplementary Table 3.** Primers used for qRT-PCR and RT-PCR.

5) **Supplementary Fig. 1.** Hit inhibitors of U2AF–RNA complexes.

6) **Supplementary Fig. 2.** Supporting evidence for *de novo* synthesis of NSC 194308.

7) **Supplementary Fig. 3.** Comparison of the anti-splicing activities of NSC 194308 variants.

8) **Supplementary Fig. 4.** Immunoblots related to Fig. 5.

9) **Supplementary Fig. 5.** Immunoblots related to Fig. 6.

**Supplementary Results: A High-Throughput Screen for U2AF–RNA Inhibitors**

To screen for inhibitors of U2AF – RNA binding (**Supplementary Table 1**), we used an *DEK(-3C)* $^{Fl}$RNA with a cytosine introduced at the -3 nucleotide relative to the spliced junction, since the S34F mutation increased binding to this splice site[37]. We chose a concentration of U2AF1$^{S34F}$–U2AF2–SF1 proteins (300 nM) near the saturated plateau of the fitted binding curve and the higher affinity $^{Fl}$*DEK(-3C)* RNA site (20 nM) (**Supplementary Fig. 1a**). Excess unlabeled *DEK(-3C)* RNA (5 μM in 1% v/v DMSO for consistency) was added as a positive control for inhibition of the complex and the effective solvent of the compounds (1% v/v DMSO) served as the negative control (for no effect). In a mock screen of positive and negative control samples, a Z′-factor [39] of 0.78 (**Supplementary Fig. 1b**) supported the expectation that the fluorescence polarization (FP) screen would identify hit inhibitors of U2AF – $^{Fl}$RNA complexes.

We screened the NCI Diversity Set V library for inhibitors of the U2AF1$^{S34F}$–U2AF2–SF1–*DEK(-3C)* $^{Fl}$RNA complex (**Supplementary Fig. 1c**), based on the premise that cancers are sensitized to spliceosome inhibition. The Z-factor of the screen was acceptable (0.53) [39]. About 5%, or 80 compounds, reduced the FP by at least 25% of that observed in the presence of excess unlabeled *DEK(-3C)* RNA (positive control). Sixteen of the initial hit compounds (1% of the starting library) reproducibly reduced the FP of the U2AF–RNA complex by >25% compared to the positive control.

Effects on the fluorescence emission or mobility of the fluorescein fluorophore could interfere with the apparent polarization and lead to a "false positive" hit. To rule out such potential artifacts, we counter-screened the sixteen compounds for alterations of the *DEK(-3C)* $^{Fl}$RNA FP in the absence of protein. Thirteen of the sixteen compounds were false hits that decreased FP during titrations of the fluorescein-labeled oligonucleotide alone. The remaining three compounds (NSC 71795, NSC 228155, NSC 207895, **Supplementary Fig. 1d**) selectively decreased FP of the U2AF1$^{S34F}$–U2AF2–SF1–*DEK(-3C)* $^{Fl}$RNA complex and had no effect on the *DEK(-3C)* $^{Fl}$RNA, suggesting that these compounds were bona fide "hits" that dissociated the U2AF protein from the RNA site. Titrations of these three hit inhibitors into the U2AF1$^{S34F}$–U2AF2–SF1–*DEK(-3C)* $^{Fl}$RNA complex showed that micromolar concentrations of two compounds reduced the FP response by half ($IC_{50}$'s approximately 8 μM for NSC 71795, 15 μM NSC 228155 and 50 μM for NSC 207895, **Supplementary Fig. 1e**).

We noticed that the most effective hit inhibitor NSC 71795 corresponds to ellipticine, a hetero-tetracyclic alkaloid that is well-known to intercalate with DNA [40,41]. Based on a common aromatic core (**Supplementary Fig. 1d**), we reasoned that NSC 228155 likewise could dissociate the U2AF–RNA complex by intercalating between the nucleic acid bases. To test this possibility, we investigated the ability of NSC 228155 to bind *DEK(-3C)* $^{Fl}$RNA in an electrophoretic mobility shift assay (**Supplementary Fig. 1f**). Addition of NSC 228155 to the RNA site produced a well-resolved band migrating above the RNA alone. The large shift in electrophoretic migration could be attributed to distortion of the RNA structure following intercalation of the compound [42]. We concluded that NSC 228155 inhibited U2AF–RNA complexes by competitively binding the RNA site. By analogy with ellipticine, the heterocyclic hit inhibitors most likely bind RNA by intercalating between the RNA bases.

**Supplementary Table 1.** Small molecule screening data for U2AF – RNA enhancers*

| Category | Parameter | Description |
|---|---|---|
| Assay | Type | Monitoring protein-RNA association |
| | Target* | U2AF1$^{S34F}$–U2AF2–SF1–*DEK(-3U)* $^{Fl}$RNA ribonucleoprotein |
| | Primary Measurement | Fluorescence polarization of 5′-fluorescein (Fl)-labeled RNA |
| | Key Reagents | Synthetic $^{Fl}$RNA oligonucleotide and purified proteins |
| | Protocol* | Protein (80 nM)–RNA (20 nM) mixture was incubated with compound (5 μM) for 60 min. Samples were excited at 490 nm and fluorescence polarization was measured at 520 nm. |
| Library | Size | 1,593 compounds |
| | Composition | Diversity Set V |
| | Source | National Cancer Institute<br>Developmental Therapeutics Program |
| Screen | Format | 384-well plates |
| | Concentration tested | 5 μM |
| | Plate controls* | Positive: 1.3 μM U2AF1$^{S34F}$–U2AF2–SF1<br>Negative: 1% v/v DMSO (matching solvent) |
| | Reagent/Dispensing system | JANUS Varispan Automated Workstation (PerkinElmer) |
| | Detection instrument | EnVision plate reader (PerkinElmer) |
| | Assay validation/QC | Z-factor 0.53; Counterscreen of $^{Fl}$RNA; Triplicate replicates; Titrations of final hit |
| | Correction factors | None |
| | Normalization | Polarization of 1% v/v DMSO control sample |
| Post-HTS Analysis | Hit criteria | Polarization increased by >75% of the signal of the positive control |
| | Hit rate* | 0.4% including false positives;<br>0.06% specifically targeting ribonucleoprotein |
| | Additional assays* | 1. Surface plasmon resonance binding<br>2. *In silico* docking<br>3. Splicing of prototypical substrate in nuclear extract<br>4. Pre-mRNA splicing in HEK 293T and K562 cell lines<br>5. Cell viability assays of HEK 293T and K562 cell lines |
| | Confirmation of hit purity and structure | Synthesized NSC 194308 with >95% purity<br>and confirmed activity |

*Conditions for the U2AF – RNA inhibitor screen were identical to the enhancer screen, apart from: *(i)* use of *DEK(-3C)* $^{Fl}$RNA oligonucleotide and higher concentration of protein (300 nM) in the assay, *(ii)* positive plate control (5 μM unlabeled RNA), *(iii)* hit criteria (polarization reduced by >25% relative to the signal of the positive control), *(iv)* hit rate (1% including all hits; 0.2% affecting ribonucleoprotein) and post-HTS analysis, which included electrophoretic mobility shift assays.

**Supplementary Table 2.** Scores for docking NSC 194308 on 3D structures.

| **Protein** | **PDB ID** | **Score** (a.u.)* |
|---|---|---|
| Open U2AF2 | 5EV4 | -35 |
| Closed U2AF2 | 2YH0 | -15 |
| SF1 KH QUA2 | 1K1G | -28 |
| SF1 coiled coil–U2AF2 UHM | 4FXW | -25 |
| U2AF1 UHM–U2AF2 ULM | 1JMT | -20 |

*Score of most favorable site in arbitrary units (a.u.). Negative values favor compound binding.

**Supplementary Table 3.** Primers used for qRT-PCR and RT-PCR.

| Gene name | Forward primer | Reverse primer |
|---|---|---|
| **Application:** | **Taqman$^{TM}$ RT-qPCR** | |
| *AdML*, probe | FAM6-AGCTGTTGGGCTGCAG-3C spacer-BHQ$^{R}$1 | |
| *AdML* | TCTCTTCCGCATCGCTGTCT | GCGAAGAGTTTGTCCTCAACGT |
| **Application:** | **qRT-PCR** | |
| *CCND1* | GGCGGAGGAGAACAAACAGA | TGTGAGGCGGTAGTAGGACA |
| *DUSP11* | GGTGTCCACTGTACCCATG | GCCTCACGCCTTCTACATCA |
| *HTATSF1* | TGTATACGTGTCTGGTTTGCC | ACAGCAAAGACCGTCTCCTT |
| *GAPDH* | TGCACCACCAACTGCTTAGC | GGCATGGACTGTGGTCATGAG |
| **Application:** | **RT-PCR** | |
| *ASUN* | GTGAAGGATGTGGAGGAAGAGTT | TCACAATAACACTGGCTAATGGA |
| *RIPK2* | CATTCCCTACCACAAACTCG | AGGATGCGAAATCTCAATGG |
| *RNF10* | GGCCCTCTCCATTTCTCCTC | TTATCACTCTCCCCGTCGCT |

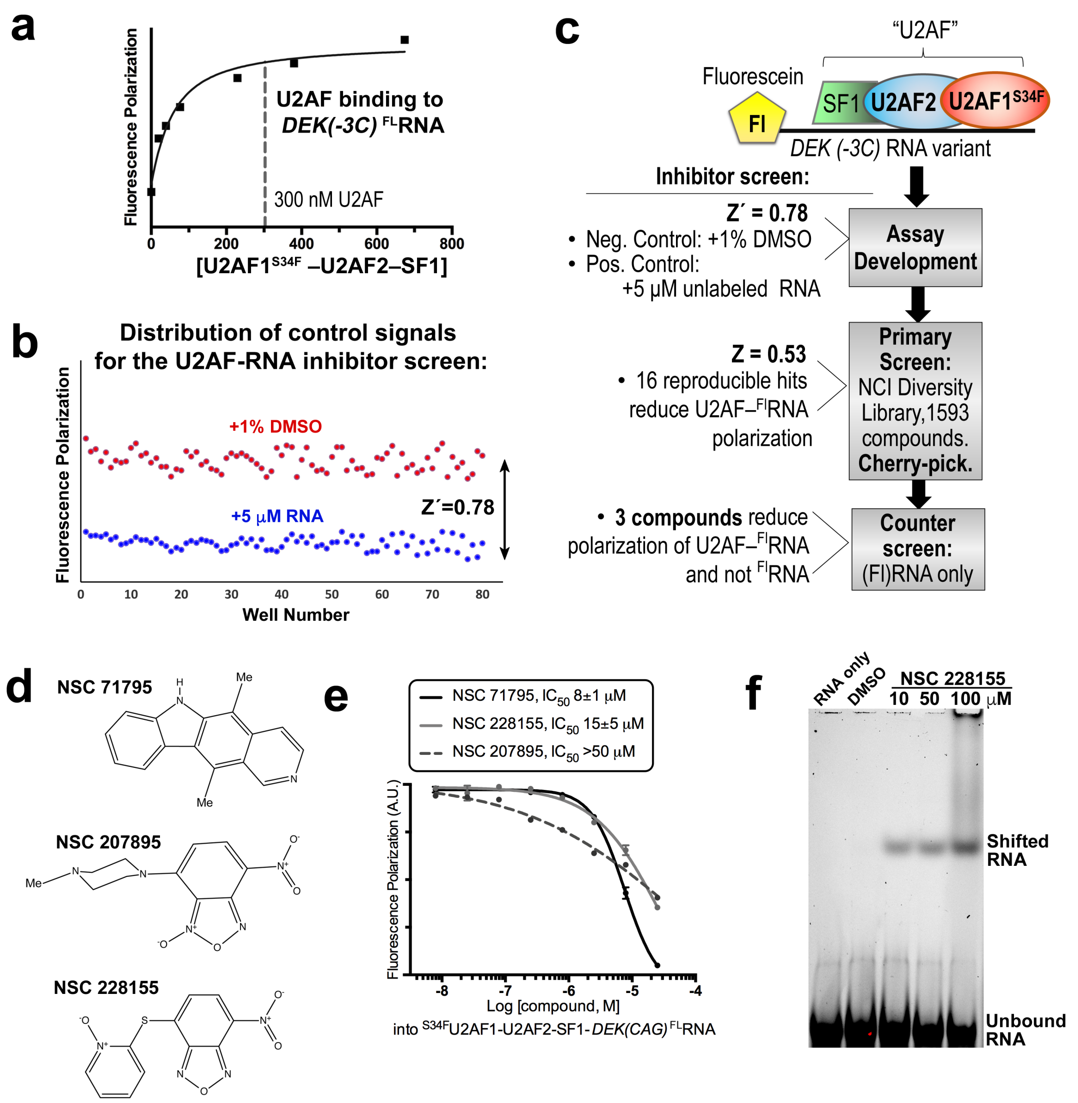

**Supplementary Fig. 1.** Hit inhibitors of U2AF–RNA complexes identified from the chemical library screen. (**a**) Fluorescence polarization binding curve for U2AF1$^{S34F}$–U2AF2–SF1 titrated into fluorescein *DEK(-3C)* $^{FL}$RNA. The nonlinear fits are overlaid on the average data points of three replicates as a solid curve. A dashed line marks the protein concentration used in each screen. (**b**) Distribution of positive (blue) and negative (red) control signals used for the calculation of the Z´-factors of the assay. (**c**) Overview of the inhibitor screen. (**d**) Chemical structures of the three hit inhibitors. Me, Methyl. (**e**) Dose-response curves of hit inhibitors titrated into U2AF1$^{S34F}$–U2AF2–SF1–*DEK(-3C)* $^{FL}$RNA (FL, 5´-fluorescein-labeled) in triplicate. The data points and error bars represent the means and standard deviations of three replicates. (**f**) Electrophoretic mobility shift assay of NSC 228155 binding *DEK(-3C)* $^{FL}$RNA. the RNA (2.5 µM) was mixed with compound at 10, 50, or 100 µM final concentrations or a 1% DMSO control in a 25 µL volume containing 8 mM $MgCl_2$, 100 mM KCl, 10 mM HEPES pH 7.4, 0.1 mg/mL BSA, 0.05% P20, 10% glycerol, 10 U Superase•In$^{TM}$ (Invitrogen by Thermo Fisher, Sci.). Following incubation for 40 min on ice, the mixture was separated on a pre-electrophoresed 5% w/v gel of 37.5:1 acrylamide/bis-acrylamide in 0.5x TBE at 4 °C and imaged following 490 nm excitation. *Related to Fig. 1.*

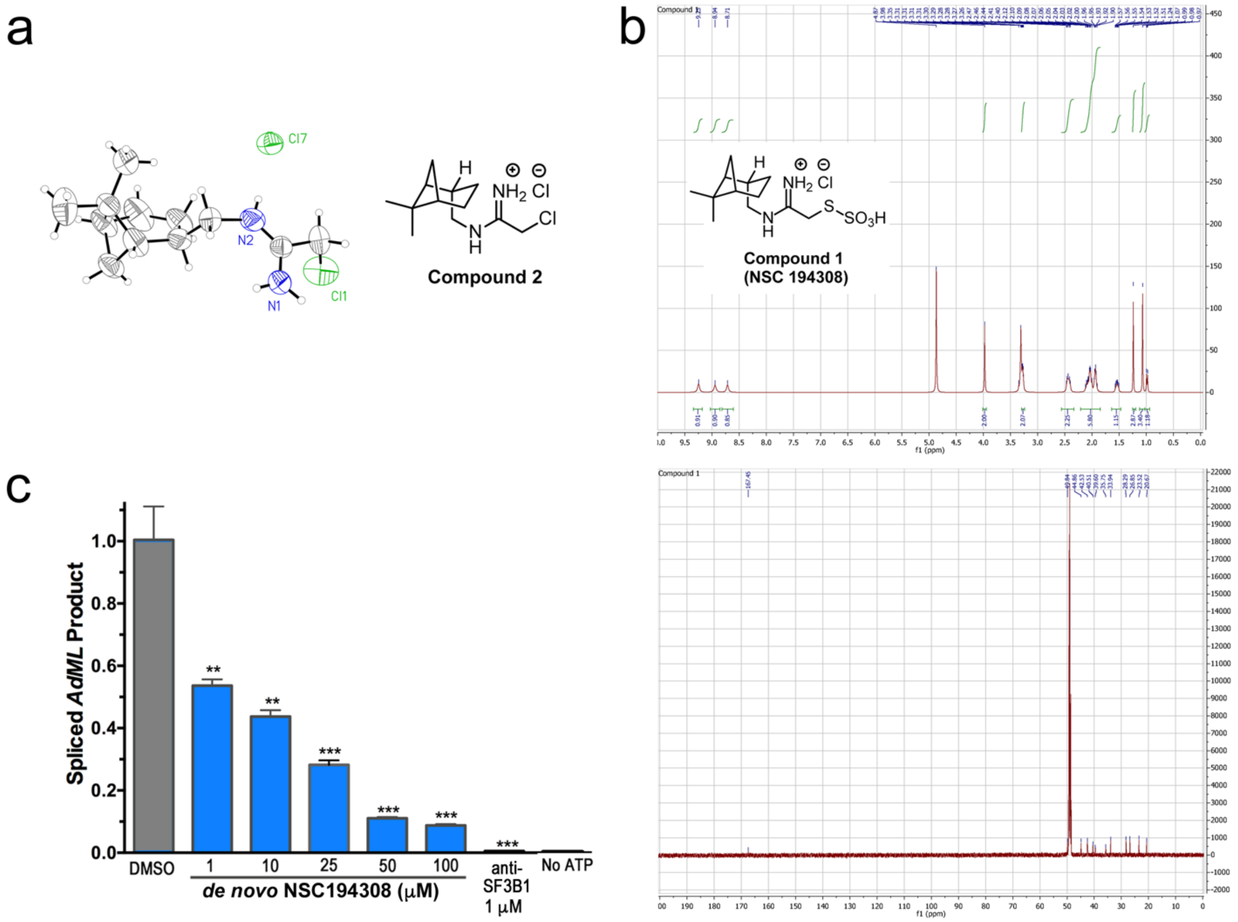

**Supplementary Fig. 2.** Supporting evidence for *de novo* synthesis of NSC 194308. (**a**) Crystal structure of Compound 2 intermediate (2-chloro-*N-(-)-cis*-myrtanyl acetamidinium hydrochloride). (**b**) NMR spectra of the final Compound 1 product (NSC 194308). (**c**) The *de novo* synthesized NSC 194308 compound inhibits *in vitro* splicing of the *AdML* substrate with similar potency as the compound obtained from the NCI DTP source (Fig. 2b). Two-tailed unpaired t-tests with Welch's correction calculated in GraphPad Prism compared the indicated NSC 194308 concentration with a 1% v/v DMSO control: **, $p$<0.005; ***, $p$<0.0005. *Related to Fig. 2.*

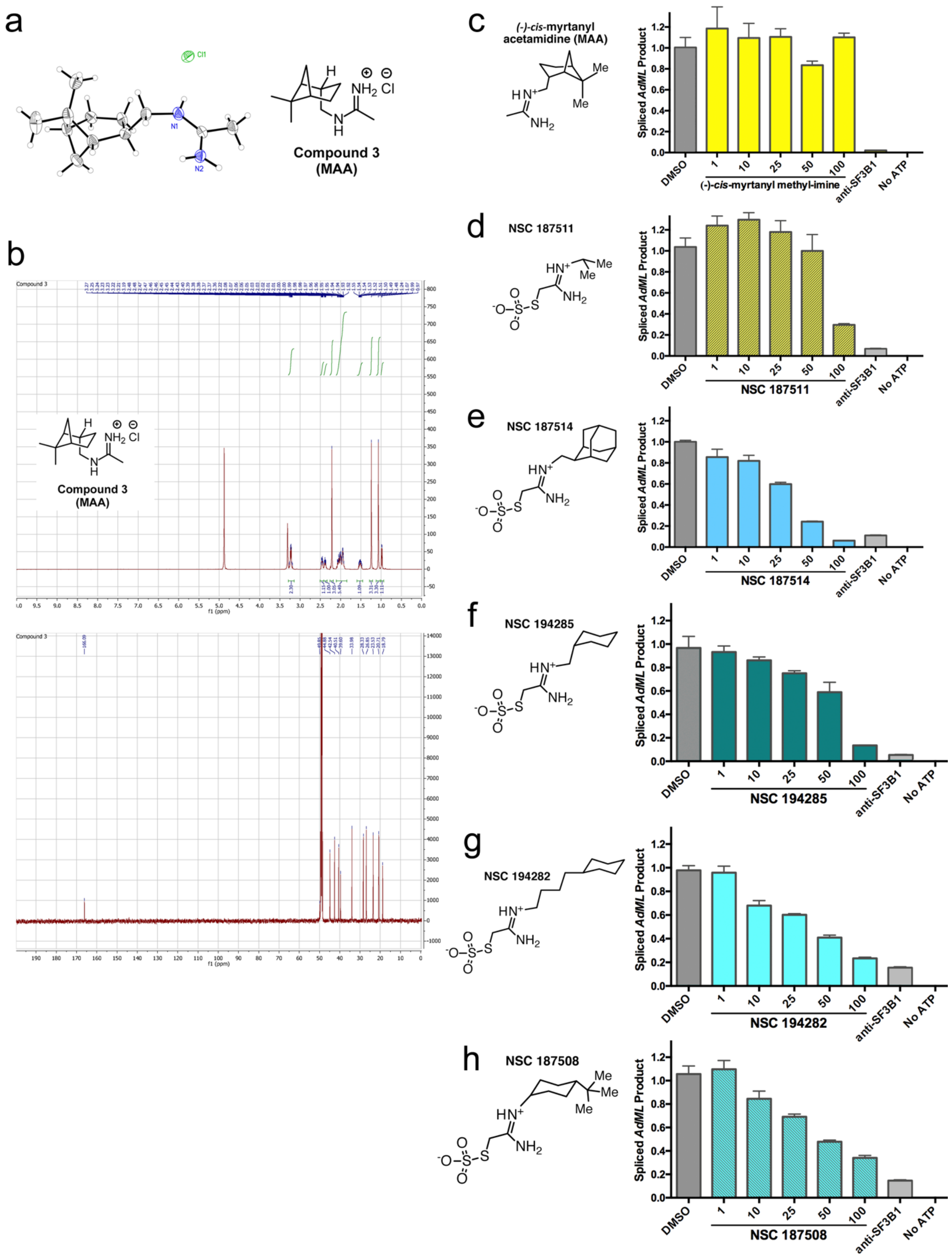

*Supplemental Fig. 3 legend cont. next page.*

**Supplementary Fig. 3.** Comparison of the anti-splicing activities of NSC 194308 variants. (**a-b**) Supporting evidence for *de novo* synthesis of **Compound 3** (*N-(-)-cis*-myrtanyl acetamidinium hydrochloride, MAA) includes the crystal structure (**a**) and NMR spectra (**b**). (**c-h**) Compounds were titrated into HeLa nuclear extract supplemented with *AdML* pre-mRNA. The spliced product was quantified by qRT-PCR and normalized to a 1% DMSO control (dark gray). The anti-SF3B1 inhibitor is pladienolide B (1 μM) (light gray). The chemical structures are shown to the left of each *in vitro* splicing result. (**c**) (-)-*cis*-myrtanyl acetamidine (MAA), (**d**) NSC 187511, (**e**) NSC 187514, (**f**) NSC 194285, (**g**) NSC 194282, (**h**) NSC 187508. MAA was synthesized *de novo* and other compounds were obtained from the NCI DTP. *Related to Fig. 4.*

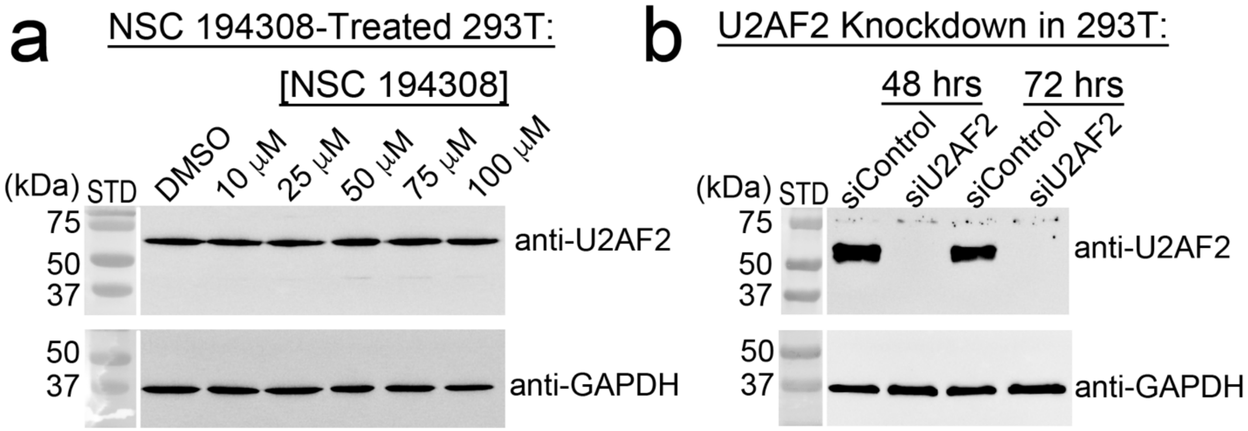

**Supplementary Fig. 4.** Immunoblots of U2AF2 levels relative to a GAPDH-loading control, for the HEK 293T samples shown in Fig. 5. (**a**) U2AF2 levels remain constant in samples taken five hours after NSC 194308 treatment. Colorimetric images of molecular weight standards (STD) are separated from the chemiluminescent blot detection by a white line. (**b**) U2AF2 levels are reduced in samples taken the indicated number of days after treatment with a U2AF2-directed Stealth$^{TM}$ siRNA, compared to a control siRNA. *Related to Fig. 5.*

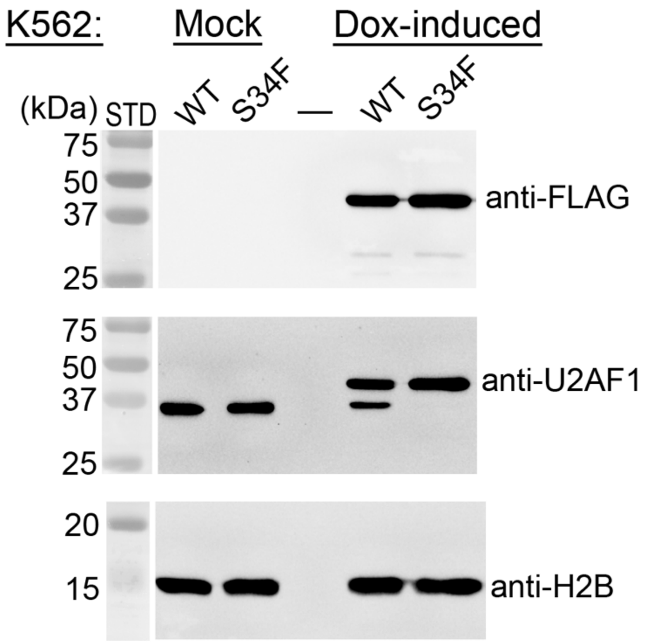

**Supplementary Fig. 5.** Immunoblots showing 3xFLAG-tagged wild-type (WT) or S34F U2AF1 expression in dox-inducible K562 cell lines. H2B, loading control. Colorimetric images of molecular weight standards (STD) are separated from the chemiluminescent detection by a white line. Doxycycline was dissolved in water immediately prior to addition at a final concentration of 250 ng $mL^{-1}$. "Mock" samples were treated in an identical manner with water replacing the doxycycline solution. *Related to Fig. 6.*
